# Semaphorin 3G (SEMA3G) is a highly selective marker of cerebral amyloid angiopathy

**DOI:** 10.64898/2026.09.10.750803

**Authors:** Kaleah Balcomb, Jessica Buchanan, Aysha Strobbe, Daphne Claus, Arline Faustin, Julie Schneider, Thomas Wisniewski, Margaret Sunde, Eleanor Drummond

## Abstract

Biomarkers of cerebral amyloid angiopathy (CAA) are critically needed. We recently identified semaphorin 3G (SEMA3G) as a novel protein selectively enriched in CAA. Here, we aimed to determine if SEMA3G was a selective marker of CAA in a large cohort of human brain tissue spanning multiple neurodegenerative diseases and three brain regions, and to determine if SEMA3G directly interacts with amyloid beta (Aβ). Multiplexed immunofluorescence showed that SEMA3G significantly accumulated only in CAA^+^ blood vessels in the brain in all cases. We also showed that SEMA3G preferentially associated with Aβ_40_, Aβ_pS8_ and Aβ_pE3,_ but not Aβ_42_. Thioflavin T assays and transmission electron microscopy showed that SEMA3G directly interacted with Aβ and slowed Aβ aggregation *in vitro*, and that this effect was more pronounced for Aβ_40_ than Aβ_42_. Together our results demonstrate that SEMA3G is a highly specific marker of CAA in the brain.

## Main

Cerebral amyloid angiopathy (CAA) is characterized by the deposition of Aβ in the walls of leptomeningeal and cortical blood vessels. CAA starts as scant Aβ deposits in the tunica media of the vessel wall near smooth muscle cells (mild CAA). As CAA severity increases, Aβ accumulates throughout the circumference of the vessel, causing progressive destruction and eventual replacement of the tunica media by Aβ deposits (moderate CAA). In some circumstances, this can lead to fragmentation of the vessel wall, resulting in microbleeds and intracerebral hemorrhages (severe CAA)^1–4^. CAA is prominent in leptomeningeal arteries, cortical arteries and arterioles and in some cases, capillaries^5^. CAA is common in Alzheimer’s disease (AD)^1–4^, where it exacerbates cognitive decline, accelerates disease progression, impairs perivascular clearance of Aβ in the brain and significantly increases risk of developing amyloid-related imaging abnormalities (ARIA) after treatment with Aβ immunotherapies^6–11^. Therefore, CAA biomarkers are urgently needed.

Currently, CAA can only be definitively diagnosed at autopsy. While several neuroimaging biomarkers for CAA have been developed, these can only provide a probable or possible diagnosis^12–14^. Identification of CAA-specific plasma or cerebrospinal fluid (CSF) biomarkers has proven challenging as most proteins enriched in CAA are also enriched in amyloid plaques, making it difficult to distinguish the two^15^. However, we recently identified several proteins selectively enriched in CAA and not in amyloid plaques by comparing the localized proteome of CAA^+^ blood vessels, CAA^−^ blood vessels, and amyloid plaques^15^. Semaphorin 3G (SEMA3G) emerged as the most promising selective marker of CAA. Both proteomics and immunostaining confirmed that SEMA3G was abundant in CAA^+^ blood vessels in both AD and mild cognitive impairment (MCI) cases, but undetectable in CAA^−^ blood vessels and amyloid plaques^15^. The highly selective enrichment of SEMA3G in CAA^+^ blood vessels is missed in bulk tissue proteomics studies of AD brain, which do not report a significant increase of SEMA3G in AD^18,19^.

Very little is known about the role of SEMA3G in CAA or AD. In the brain, SEMA3G is almost exclusively expressed by arterial endothelial cells^16,17^. Expression is low in capillary endothelial cells, and no expression is observed elsewhere in the brain, including in venous endothelial cells^16,17^. Functionally, SEMA3G is a secreted protein that acts as a signaling molecule to a variety of cell types via complexing with neuropilin 2/plexins to regulate angiogenesis, smooth muscle cell proliferation and synapse formation^20–26^.

This study aimed to significantly extend our previous results to determine: (1) if SEMA3G was consistently a selective marker of CAA in a large, diverse cohort of human cases, across three brain regions with varying CAA severity, and in the context of variety of neurodegenerative diseases (AD, progressive supranuclear palsy, corticobasal degeneration, Pick’s disease, Lewy body dementia, no neurodegenerative disease), and (2) if SEMA3G directly interacted with specific Aβ species present in CAA.

## Results

### SEMA3G is a marker of CAA and not amyloid plaque pathology

To characterise SEMA3G in the brain, immunohistochemistry was initially performed on inferior temporal cortex sections from cases of mild CAA (*n* = 14), moderate CAA (*n* = 16), moderate CAA without Aβ plaque pathology (CAA^+^ P^−^, *n* = 5), Aβ plaque pathology without CAA (CAA^−^ P^+^, *n* = 20) and age- and sex-matched controls without CAA or Aβ plaques (*n* = 20) (Supplementary Table 1). Machine learning was used to segregate and measure Aβ plaques, CAA^+^ and CAA^−^ vessels in the grey matter based on immunostaining of Aβ and the vascular marker COL4A1 (Extended Data Fig. 1), and SEMA3G immunofluorescence assessed in each.

Strikingly, SEMA3G consistently localised to CAA^+^ vessels and was not observed in Aβ plaques (Fig. 1a). SEMA3G immunoreactivity in blood vessels showed a strong positive correlation to CAA load (Spearman’s ⍴ = 0.523, *p* < 0.0001) but not plaque load (Spearman’s ⍴ = 0.018, *p* = 0.88) (Fig. 1b, c). Negligible levels of SEMA3G immunofluorescence (0.06±0.2%) colocalised with amyloid plaques (*n* = 38), which was consistent across cases with and without CAA co-pathology (Fig. 1d). This was particularly notable in instances of dyshoric CAA, where SEMA3G only colocalized with the vascular CAA component and not in the plaque-like Aβ deposit surrounding the vessel (Fig. 1e). Quantification of total SEMA3G levels in the vasculature revealed that SEMA3G levels were high in moderate CAA (0.50±0.43% of vessel area) and CAA^+^ P^−^ cases (0.28±0.24%) (Fig. 1f). In contrast, SEMA3G levels were low in controls (0.08±0.12%), CAA^−^ P^+^ (0.06±0.08%) and mild CAA (0.07±0.07%). In all cases, SEMA3G was almost exclusively localized to blood vessels (98.07±2.57% of total signal across all cases) and was primarily observed in larger vessels in the grey and white matter, not capillaries (Fig. 1g).

**Figure 1:**
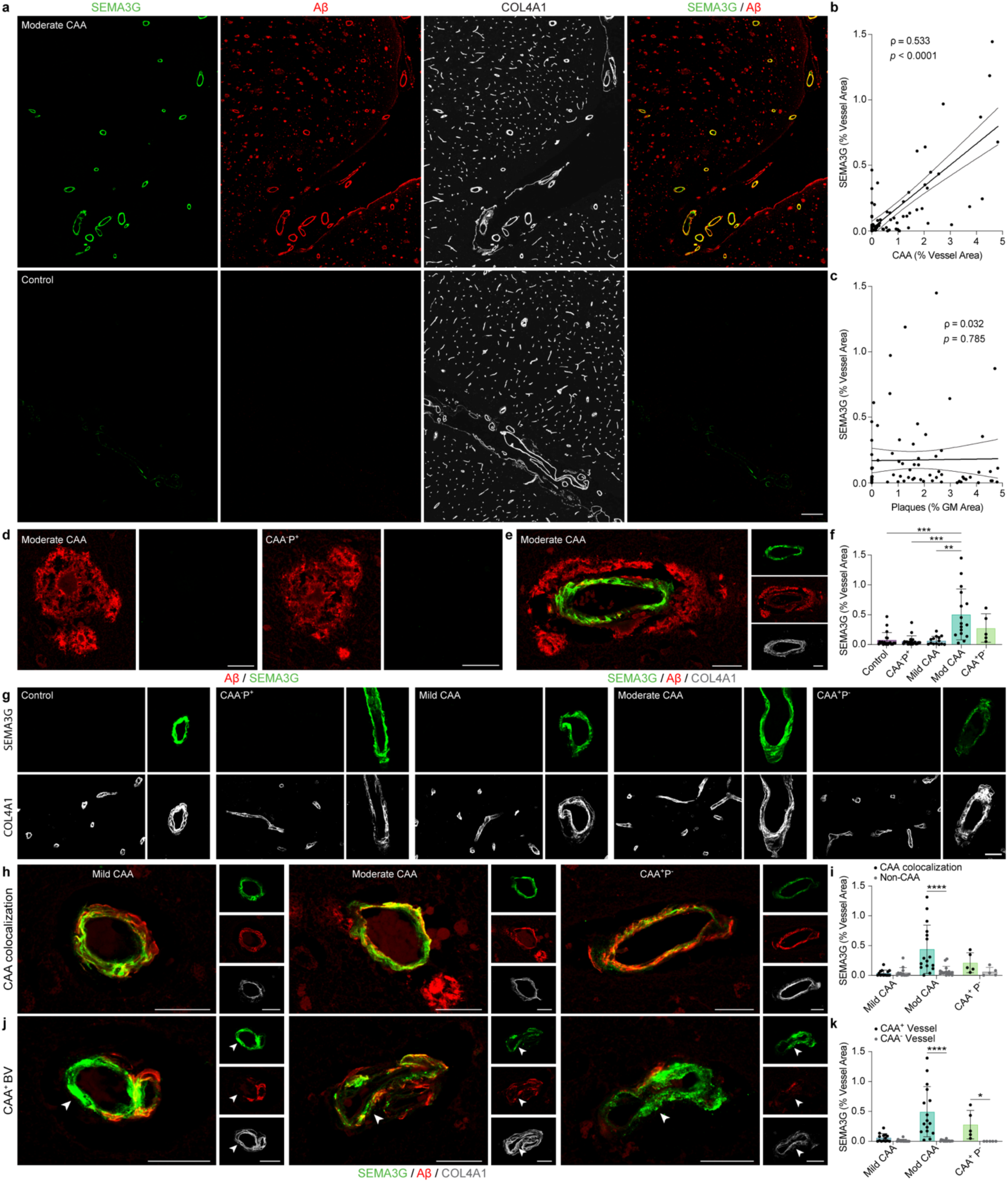
SEMA3G is strongly increased in CAA vessels in the inferior temporal cortex. **a,** SEMA3G immunofluorescence colocalized with CAA and not plaques in cases of AD with CAA, with minimal signal in controls. **b-c,** SEMA3G was strongly correlated to CAA load **(b)** but not amyloid plaque load **(c)**. Nonparametric Spearman correlations of *n* = 75, with 95% confidence intervals. **d,** Representative images showing an absence of SEMA3G in amyloid plaques in cases with and without CAA. **e,** Representative image of a CAA-positive vessel surrounded by a dyshoric plaque, showing SEMA3G only in the vessel. **f,** SEMA3G signal in the grey matter was significantly increased in Moderate CAA cases compared to controls, CAA^−^ P^+^ and Mild CAA. Data show mean ± SD of *n* = 20 control, CAA^−^ P^+^, *n* = 14 Mild CAA, *n* = 16 Moderate CAA, *n* = 5 CAA^+^ P^−^; \**p*<0.05, \*\**p*<0.01, \*\*\**p*<0.001, determined by a Kruskal-Wallis test with Dunn’s post-hoc analysis. **g,** SEMA3G immunofluorescence was almost exclusively observed in large blood vessels and not capillaries, regardless of amyloid pathology type or abundance. **h,** SEMA3G colocalization with CAA was evident in all CAA-positive cases, irrespective of severity. **i,** Quantification of SEMA3G colocalization with CAA. **j,** SEMA3G immunofluorescence was observed throughout CAA^+^ vessels in all conditions. Arrows indicate areas of SEMA3G immunofluorescence independent of Aβ. **k,** Quantification of SEMA3G localization to CAA^+^ and CAA^−^ vessels. Data in i, k show mean ± SD of *n* = 14 Mild CAA, *n* = 16 Moderate CAA, *n* = 5 CAA^+^ P^−^; \**p*<0.05, \*\*\*\**p*<0.0001, determined by two-way ANOVA with Tukey’s post-hoc. Scale bar = 200µm (a), 25 µm (d,e,g,h,j).

### SEMA3G strongly localizes to CAA^+^ blood vessels

Given the strong localization of SEMA3G to blood vessels, and the increase of SEMA3G signal in CAA-positive cases, we next examined the localization pattern of SEMA3G in CAA*^+^*vessels in cases with differing CAA severity. SEMA3G showed increased colocalization with Aβ in CAA as CAA severity increased; 53.9±32.6% of SEMA3G immunoreactivity colocalized with Aβ in mild CAA, 77.5±23.3% in moderate CAA, and 82.5±11.7% in CAA^+^ P^−^ cases (Fig. 1h, i). Interestingly, we observed that SEMA3G signal in CAA*^+^* vessels was not limited to regions with Aβ deposition, but was instead broadly increased throughout the circumference of CAA*^+^* vessels. We therefore investigated the levels of SEMA3G immunofluorescence in CAA*^+^*versus CAA^−^ blood vessels. CAA^+^ vessels were defined by presence of any Aβ, including trace, punctate deposits. Remarkably, 98.9±0.9% and 92.7±9.6% of SEMA3G immunoreactivity was present in CAA*^+^* vessels in CAA^+^ P^−^ and moderate CAA cases respectively (Fig. 1j, k). In mild CAA cases, 80.3±29.0% of SEMA3G immunoreactivity was found in CAA*^+^* vessels, showing a clear accumulation of SEMA3G in blood vessels with CAA even in the earliest stage of CAA development.

### SEMA3G immunofluorescence shows consistent trends between temporal and occipital cortices

To investigate if SEMA3G similarly localized to CAA^+^ vessels in a different brain region, *n* = 4 moderate CAA, *n* = 6 mild CAA, *n* = 4 CAA^+^ P^−^, *n* = 4 CAA^−^ P^+^ and *n* = 5 control cases from the primary visual cortex were assessed. CAA load characterisation showed similar trends to the inferior temporal cortex, while plaque load was slightly lower in the primary visual cortex compared to the inferior temporal cortex (*p <* 0.05 in mild CAA only) (Extended Data Fig. 2).

In the primary visual cortex, SEMA3G showed a similar staining pattern to that observed in the interior temporal cortex. Basal levels of SEMA3G were low and were similar between controls (0.002±0.002% of blood vessel area), CAA^−^ P^+^ (0.03±0.03%) and mild CAA (0.04±0.04%). SEMA3G was significantly increased in blood vessels in both moderate CAA (0.17±0.09%, *p* < 0.01) and CAA^+^ P^−^ (0.12±0.11%, *p* < 0.05) compared to controls (Fig. 2a). SEMA3G levels in the primary visual cortex vasculature again had a strong correlation to CAA load (Spearman’s ⍴ = 0.822, *p* < 0.0001) (Fig. 2b) but not plaque load (Spearman’s ⍴ = 0.213, *p* = 0.33) (Fig. 2c). 98.6±1.7% of SEMA3G immunofluorescence in the primary visual cortex localized to blood vessels, and was again primarily observed in larger blood vessels across all cases (Fig. 2d). Negligible SEMA3G was observed in plaques (0.005±0.02% of total SEMA3G signal, *n* = 12 plaque-positive cases), with dyshoric plaques around CAA*^+^* vessels similarly negative for SEMA3G (Fig. 2e). Colocalization of SEMA3G with Aβ in CAA occurred at similar proportions to that observed in the inferior temporal cortex; 44.6±29.8% of SEMA3G immunoreactivity colocalized with Aβ in CAA in mild CAA cases, 75.2±26.0% in moderate CAA, and 89.7±6.1% in CAA^+^ P^−^ (Fig. 2f). We again observed that SEMA3G accumulated throughout CAA*^+^* vessels, including regions that extended beyond the Aβ deposition; 72.1±21.9% in mild CAA, 84.3±17.0% in moderate CAA and 98.4±1.3% in CAA^+^ P^−^ (Fig. 2g,h).

**Figure 2:**
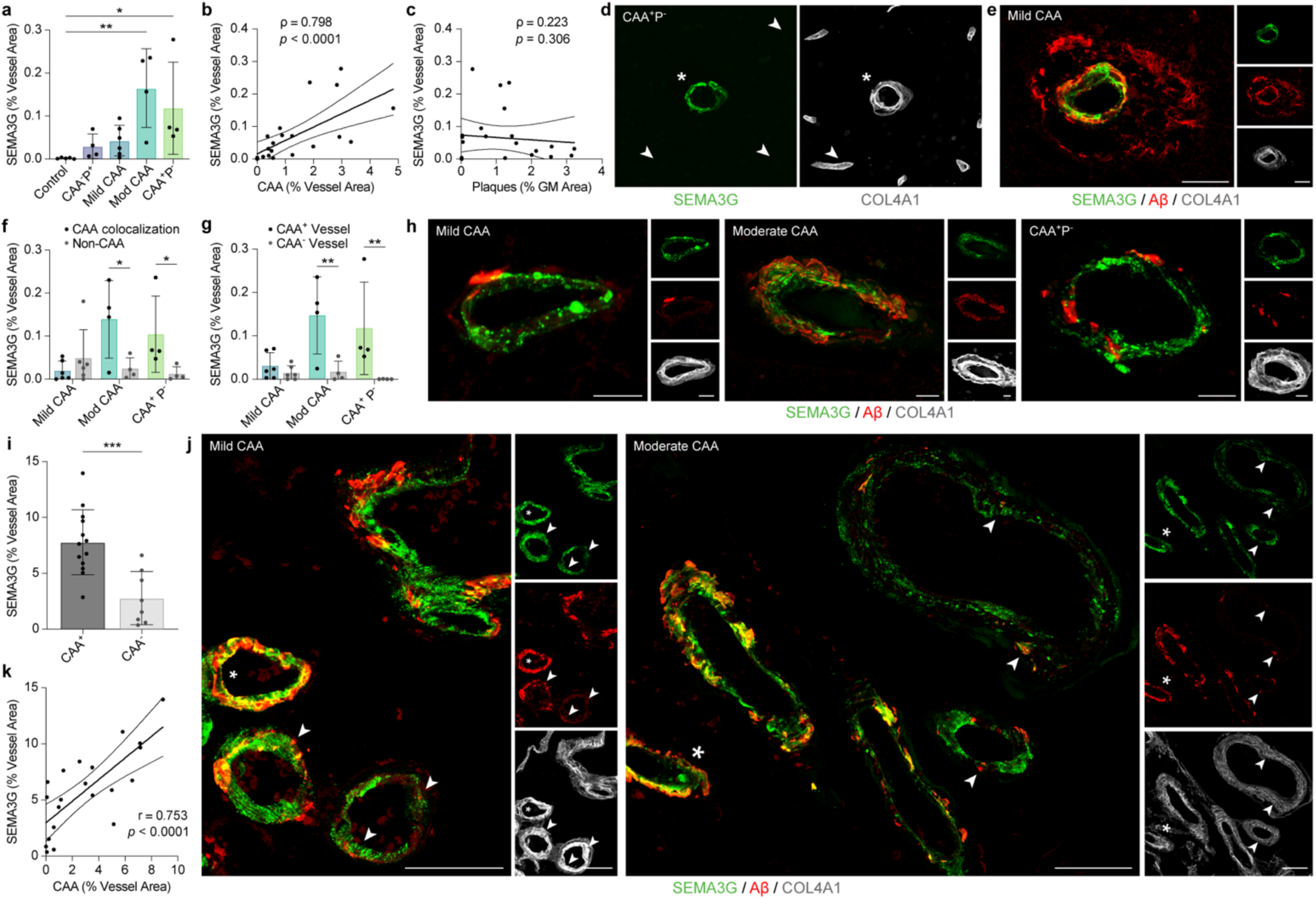
SEMA3G is strongly increased in CAA vessels in the primary visual cortex. **a,** SEMA3G immunoreactivity was significantly increased in Moderate CAA and CAA^+^ P^−^ compared to controls. Data show mean ± SD of *n* = 5 control, *n* = 6 Mild CAA, *n* = 4 Moderate CAA, CAA^−^ P^+^, CAA^+^ P^−^; \**p* < 0.05, \*\**p* < 0.01 determined by a Kruskal-Wallis test with Dunn’s post-hoc analysis. **b-c,** SEMA3G immunofluorescence was strongly correlated to CAA load **(b)** but not amyloid plaque load **(c).** Nonparametric Spearman correlation of *n* = 23 cases with 95% confidence intervals. **d,** Representative image showing SEMA3G localization to large blood vessels (star), and not small blood vessels (arrows). **e,** Representative image showing the absence of SEMA3G in dyshoric plaques around CAA^+^ vessels. **f,** The majority of SEMA3G immunofluorescence colocalized with CAA in Moderate CAA and CAA^+^ P^−^ cases. **g-h,** SEMA3G immunofluorescence was observed throughout CAA^+^ vessels in all conditions. Data in f,g show mean ± SD of *n* = 6 Mild CAA, *n* = 4 Moderate CAA, CAA^+^ P^−^; \**p*<0.05, \*\**p*<0.01, determined by two-way ANOVA with Tukey’s post-hoc. **i,** SEMA3G was increased in meningeal blood vessels of CAA^+^ cases compared to CAA^−^. Data show mean ± SD of *n* = 8 CAA^−^, *n* = 13 CAA^+^ cases; \*\*\**p* < 0.001, determined by Welch’s t test. **j,** SEMA3G strongly localized to meningeal vessels with low (arrows) or high (star) levels of CAA. **k,** SEMA3G was strongly correlated to CAA load in the meninges. Pearson correlation of *n =* 21 cases, dotted lines represent 95% confidence intervals. Scale bars = 25 µm (d,e), 10 µm (h), 50 µm (j).

### SEMA3G is elevated in CAA^+^ leptomeningeal blood vessels

Due to the high abundance of meningeal vessels present in primary visual cortex sections, we were also able to assess SEMA3G levels in the leptomeninges in this region. For this analysis, cases were reclassified into CAA^+^ (*n* = 13) or CAA^−^ (*n* = 8) based solely on leptomeningeal CAA load (Extended Data Fig. 2). SEMA3G was significantly increased in the leptomeninges of CAA^+^ cases (7.79±2.9%) compared to CAA^−^ cases (2.8±2.4%, *p* < 0.001) (Fig. 2i). In a similar trend to cortical vessels with CAA, SEMA3G preferentially accumulated in CAA*^+^* vessels in the leptomeninges (82.8±20.2% of SEMA3G immunoreactivity) (Fig. 2j, Extended Data Fig. 2). This was despite only 18.1±11.4% of SEMA3G immunoreactivity in leptomeningeal vessels directly colocalizing with Aβ in CAA (Extended Data Fig. 2). SEMA3G levels in the leptomeninges again showed a strong correlation to CAA Load (Pearson’s r = 0.753, *p* < 0.0001) (Fig. 2k).

### SEMA3G immunoreactivity closely reflects that of Aβ40 in vessels throughout the parenchyma and leptomeninges

We were next interested to determine if SEMA3G preferentially colocalized with a specific Aβ proteoform. Colocalization of SEMA3G with Aβ_40_, Aβ_42_, pyroglutamated Aβ (Aβ_pE3_) and Aβ phosphorylated at serine 8 (Aβ_pS8_) was assessed using immunofluorescence on *n* = 5 primary visual cortex sections with a range of pathology (*n* = 2 mild CAA, *n* = 2 moderate CAA, *n* = 1 CAA^+^ P^−^). Serial sections were used where possible (Supplementary Table 1).

SEMA3G immunofluorescence most closely paralleled Aβ_40_ in the vasculature of all cases. In mild CAA cases, large parenchymal vessels with strong SEMA3G immunofluorescence were most commonly positive for Aβ_40_, but negative for Aβ_42_, Aβ_pE3_ and Aβ_pS8_ (Fig. 3a), with occasional vessels observed to be positive for all four Aβ proteoforms. Leptomeningeal vessels also showed a similar pattern; cases with a lower meningeal CAA load had strong SEMA3G and Aβ_40_ signal, but were negative for Aβ_42_, Aβ_pE3_ and Aβ_pS8_ (Fig. 3b). Moderate CAA cases frequently showed Aβ_40_, Aβ_42_, Aβ_pE3_ and Aβ_pS8_ signal in SEMA3G-positive vessels throughout the parenchyma (Fig. 3c) and leptomeninges (Fig. 3d).

**Figure 3:**
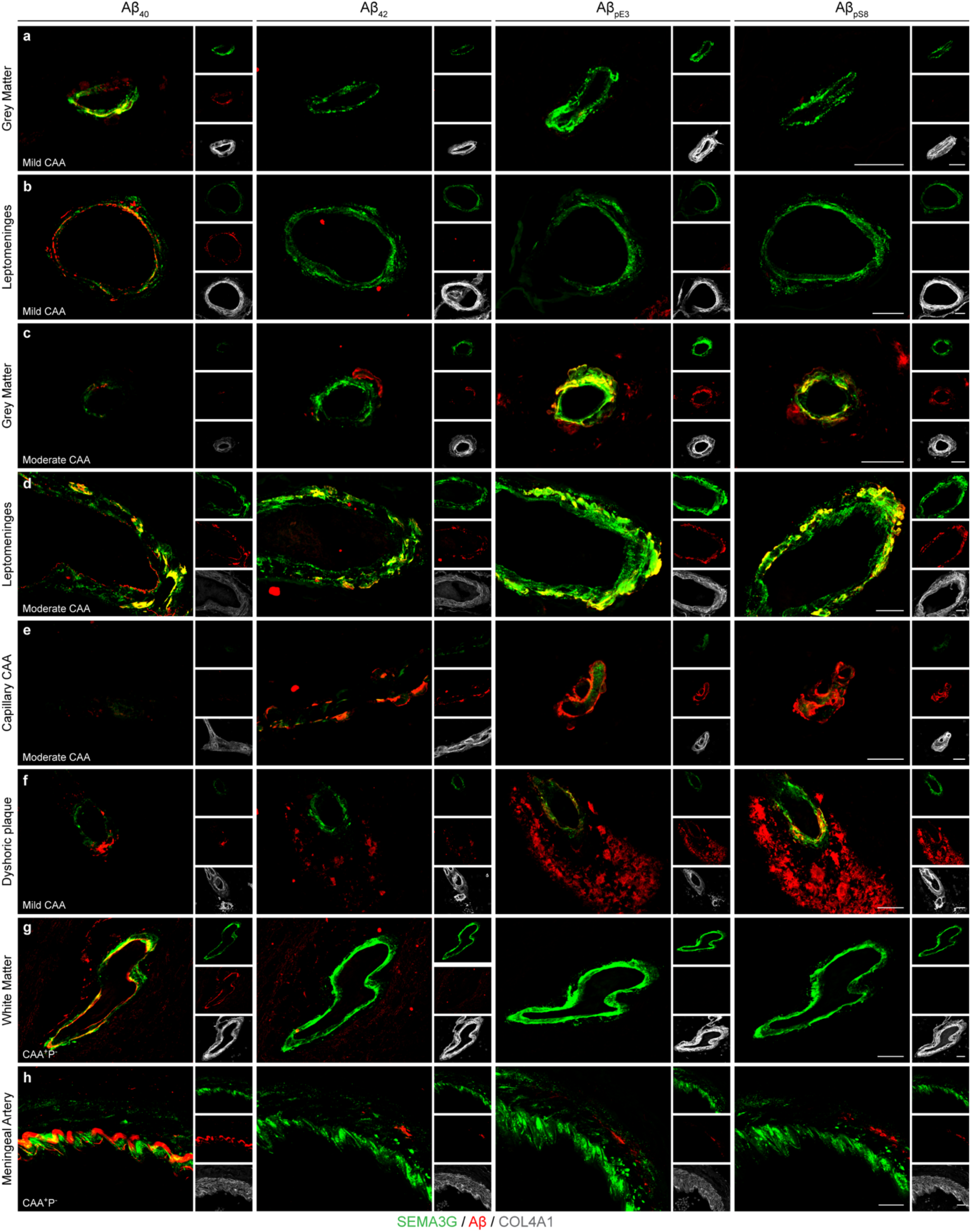
SEMA3G levels reflect Aβ40 abundance in multiple vessel types throughout the parenchyma and leptomeninges. Representative images showing immunofluorescence of SEMA3G with Aβ proteoforms throughout the parenchyma and leptomeninges. **a-b,** Mild CAA cases frequently showed SEMA3G signal in vessels with Aβ40 but not Aβ42, AβpE3 or AβpS8 in both the grey matter **(a)** and leptomeninges **(b)**. **c-d,** Moderate CAA cases predominantly showed SEMA3G immunofluorescence in vessels with Aβ40, Aβ42, AβpE3 and AβpS8 in both the grey matter **(c)** and leptomeninges **(d)**. **e,** Minimal signal from SEMA3G or Aβ40 was observed in capillary CAA. **f,** Dyshoric plaques surrounding CAA^+^ vessels were negative for SEMA3G and Aβ40. **g,** White matter vessels with strong SEMA3G and Aβ40 signal were negative for Aβ42, AβpE3 or AβpS8. **h,** Large meningeal arteries showed strong SEMA3G and Aβ40 colocalization in the endothelium in all cases. Images are representative of primary visual cortex sections from mild and moderate CAA and CAA^+^ P^−^. Scale bars = 25 µm.

Capillary CAA showed strong Aβ_42_, Aβ_pE3_ and Aβ_pS8_ immunoreactivity, but only low levels of Aβ_40_ and SEMA3G (Fig. 3e). Similarly, dyshoric plaques surrounding CAA^+^ vessels were negative for SEMA3G and Aβ_40_, but showed strong Aβ_42_, Aβ_pE3_ and Aβ_pS8_ (Fig. 3f). Interestingly, all cases exhibited large white matter vessels with strong SEMA3G and Aβ_40_ immunofluorescence, which were negative for Aβ_42_, Aβ_pE3_ or Aβ_pS8_ (Fig. 3g). Large meningeal arteries showed distinct colocalization of SEMA3G with Aβ_40_ in the endothelium, which was not observed with Aβ_42_, Aβ_pE3_ or Aβ_pS8_ (Fig. 3h). Despite the similarity observed between SEMA3G and Aβ_40_ in the vasculature, SEMA3G was not present in Aβ_40_-positive amyloid plaques (Extended Data Fig. 3), further highlighting the specificity of SEMA3G for CAA.

During our investigation of SEMA3G colocalization with Aβ proteoforms, we made the unexpected observation that our pan-Aβ antibody did not detect all Aβ_40_ pathology, including some Aβ_40_-specific CAA (Extended Data Fig. 4). We therefore hypothesized that some vessels classified as CAA^−^ using our pan-Aβ antibody may be CAA*^+^*based on the more sensitive Aβ_40_ staining. To test this, we stained a control inferior temporal cortex section (Control 1) which had high SEMA3G immunoreactivity (0.467% of blood vessel area) for SEMA3G, Aβ_40_ and pan-Aβ. We discovered that almost all vessels with SEMA3G immunofluorescence were immunoreactive for Aβ_40_ (Fig 4), which was observed consistently in both the parenchyma and meninges. In the parenchyma, even vessels with trace levels of Aβ_40_ exhibited strong SEMA3G immunofluorescence, and SEMA3G signal remained high as Aβ_40_ deposition increased (Fig. 4a). Both SEMA3G and Aβ_40_ showed strong immunofluorescence in the leptomeninges, particularly in large meningeal arteries (Fig. 4b). These results clearly demonstrate the specificity of SEMA3G as a highly selective marker of CAA, based on the ability to detect CAA*^+^* vessels to a greater level than a pan-Aβ antibody.

**Figure 4:**
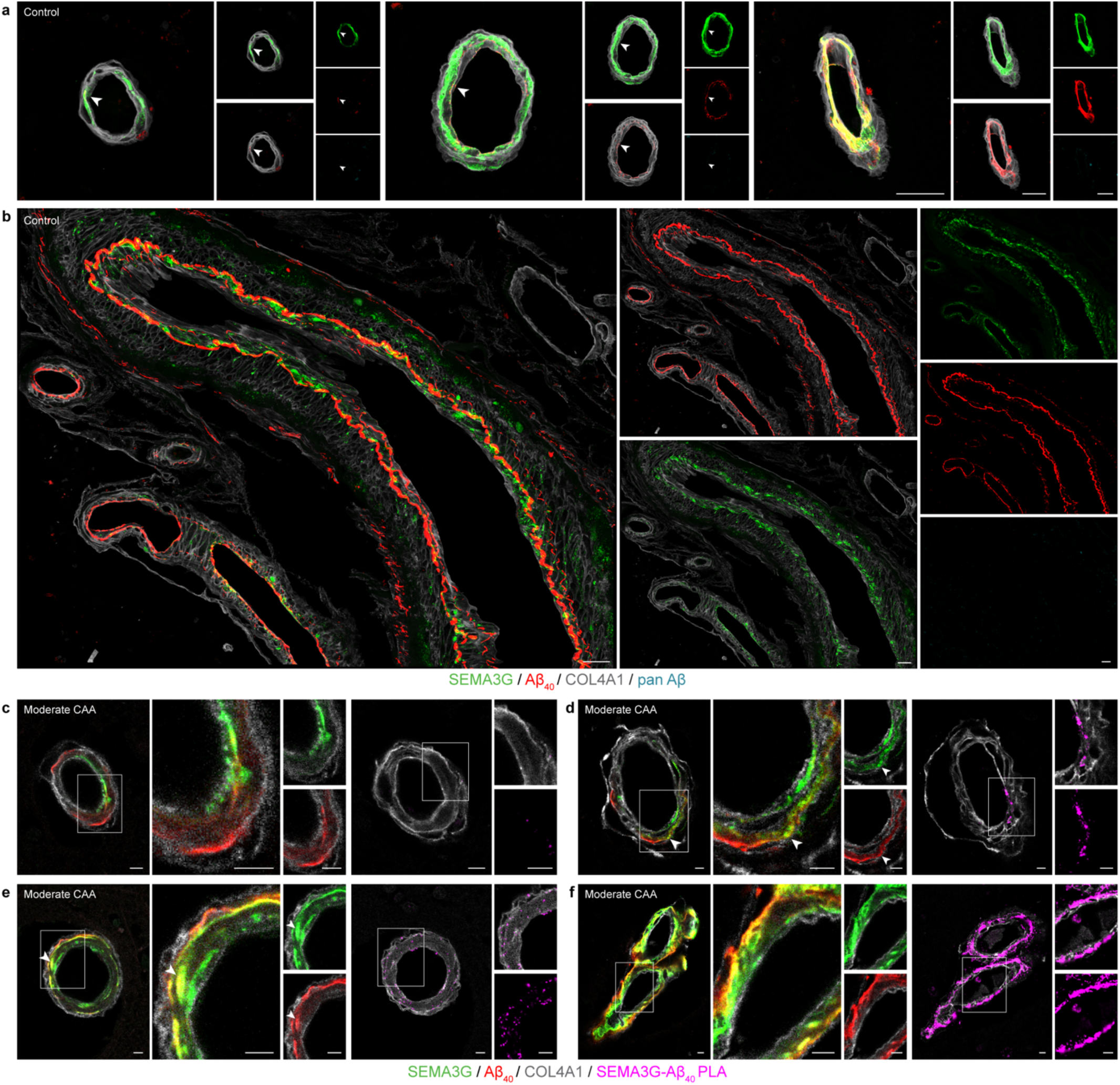
SEMA3G is consistently observed in Aβ40-positive vessels. **a-b,** Representative images of vessels from a control case showing Aβ40 immunofluorescence in the absence of pan-Aβ. **a,** SEMA3G immunofluorescence was strong in parenchymal vessels exhibiting a range of Aβ40 intensities. Arrows indicate areas with trace levels of Aβ40. **b,** Leptomeningeal vessels showed high levels of both SEMA3G and Aβ40. **c-f,** Representative single plane images of SEMA3G and Aβ40 immunofluorescence, with corresponding SEMA3G-Aβ40 proximity ligation assay (PLA) signal from a serial section. **c,** SEMA3G and Aβ40 signal was frequently observed in the same vessel but remained largely independent. PLA signal in these vessels was negligible. **d-e,** Vessels with moderate SEMA3G and Aβ40 showed some immunofluorescent colocalization (arrows), and low level PLA signal. **f,** High abundance of both SEMA3G and Aβ40 corresponded with a strong PLA signal. Images are representative of *n* = 3 moderate CAA primary visual cortex sections. Scale bars = 25 µm (a, b), 5µm (c-f).

We then performed proximity ligation assays (PLA) to more closely examine colocalization and to assess the likelihood of a potential interaction between SEMA3G and Aβ proteoforms. This approach identifies proteins that are located within 40 nm of each other. We discovered that parenchymal vessels with low levels of Aβ_40_ but strong SEMA3G exhibited minimal PLA signal. Examination of single plane confocal images revealed that while Aβ_40_ and SEMA3G were present in the same vessel, signal was found to be largely discrete (Fig. 4c). Interestingly, as immunofluorescent signal of Aβ_40_ increased and apparent colocalization with SEMA3G was abundant, PLA signal remained low (Fig. 4d-e). Vessels with high abundance of both Aβ_40_ and SEMA3G displayed strong PLA signal (Fig. 4f).

### SEMA3G is in close proximity to AβpE3 and AβpS8

To investigate if a similar relationship was occurring with Aβ_42_, Aβ_pE3_ and Aβ_pS8_, PLA for SEMA3G and Aβ proteoforms was performed on sections from *n* = 3 moderate CAA cases (Supplementary Table 1) with immunofluorescence performed on serial sections as a colocalization reference.

In the parenchyma, vessels with low levels of Aβ_42_, Aβ_pE3_ and Aβ_pS8_ immunofluorescence showed no PLA signal for Aβ_42_, but featured very strong PLA signal for both Aβ_pE3_ and Aβ_pS8_ (Fig. 5a). PLA signal for Aβ_pE3_ and Aβ_pS8_ increased as Aβ abundance in vessels increased (Fig. 5b). In contrast, PLA signal for SEMA3G-Aβ_42_ remained minimal, even in cases with Aβ_42_-positive CAA (Fig. 5b). Leptomeningeal vessels showed a similar trend to the parenchymal vessels, with strong PLA signal apparent for Aβ_pE3_ and Aβ_pS8_ (Fig. 5c). Vessel coverage and intensity of PLA signal intensity increased for Aβ_40_, Aβ_pE3_ and Aβ_pS8_ as Aβ immunoreactivity increased (Fig. 5d). Interestingly, even when Aβ_42_ was relatively abundant, PLA signal was markedly lower than that observed for the other three proteoforms (Fig. 5d). No PLA signal was observed for any Aβ proteoform in capillary CAA, reinforcing initial immunofluorescence observations (Extended Data Fig. 5).

**Figure 5:**
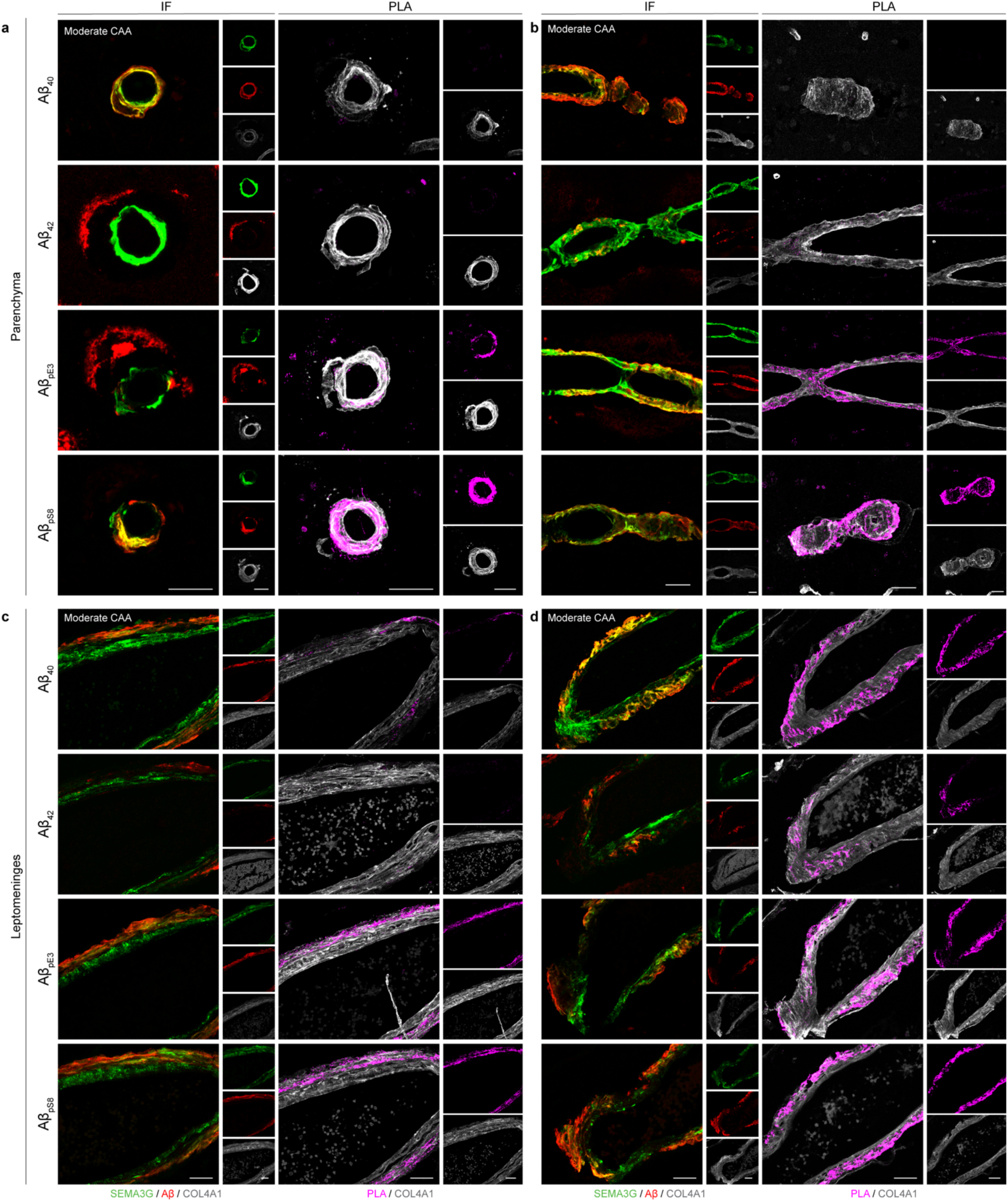
Proximity ligation assay results for SEMA3G with Aβ40, Aβ42, AβpE3 and AβpS8. Representative images of SEMA3G and Aβ proteoform immunofluorescence, with PLA performed on serial sections. **a,** Parenchymal vessel with low incidence of Aβ40, Aβ42, AβpE3 and AβpS8 immunofluorescence, and strong PLA for AβpE3 and AβpS8. **b**, Parenchymal vessel with moderate Aβ42 and high Aβ40, AβpE3 and AβpS8 immunofluorescence, with strong PLA for AβpE3 and AβpS8. **c,** Meningeal vessel with low incidence of Aβ42, moderate Aβ40, AβpE3 and AβpS8 immunofluorescence, showing low Aβ40 PLA and strong PLA for AβpE3 and AβpS8. **d,** Meningeal vessel with moderate Aβ42 and high Aβ40, AβpE3 and AβpS8 immunofluorescence, with strong PLA for Aβ40, AβpE3 and AβpS8. Images are representative of *n* = 3 moderate CAA primary visual cortex sections. Scale bars = 25 µm.

### SEMA3G is a marker of CAA regardless of co-pathology

Given the strong relationship of SEMA3G with CAA independent of Aβ plaque pathology observed in AD, we hypothesized that SEMA3G would also be a consistent marker of CAA in the context of other neurodegenerative diseases. To investigate this, immunohistochemistry was performed on superior frontal cortex sections from cases of primary and secondary tauopathies; AD + CAA (*n* = 6), Pick’s Disease (PiD) + CAA (*n* = 2), Progressive Supranuclear Palsy (PSP) + CAA (*n* = 4), Corticobasal Degeneration (CBD) + CAA (*n* = 1), or cases with aggregated alpha-synuclein; Lewy Body Disease (LBD) + CAA (*n* = 2). CAA-negative cases were also included to assess baseline SEMA3G immunofluorescence in these diseases (*n* = 4 AD, *n* = 5 PiD, PSP, CBD, LBD) alongside controls (*n* = 5). Due to the enhanced specificity of Aβ_40_ for CAA detection in our prior study, Aβ_40_ was used in this cohort to identify CAA. Assessment was limited to parenchyma due to limited presence of leptomeninges in the sections used in this experiment.

Consistent with our prior results, SEMA3G consistently and specifically only accumulated in CAA*^+^* vessels, irrespective of neurodegenerative disease context (Fig. 6). Consistent with results from the inferior temporal and primary visual cortices, SEMA3G immunofluorescence in the superior frontal cortex in AD cases was strong in CAA*^+^* vessels (Fig. 6a) and was absent in amyloid plaques (Extended Data Fig. 6). SEMA3G was unaffected by aggregated tau co-pathologies, showing strong signal in CAA^+^ vessels of PiD and PSP (Fig. 6b, c), with minimal signal observed elsewhere. Only 1 case of CBD + CAA was available for assessment, which had sparse CAA. Despite this, SEMA3G still accumulated in the few CAA*^+^* vessels present (Fig. 6d). Abundant SEMA3G was also observed in CAA*^+^* vessels in LBD (Fig. 6e). Minimal SEMA3G immunofluorescence was observed in controls (Fig. 6f) or CAA-negative PSP, CBD, PiD, LBD and AD cases (Extended Data Fig. 6), indicating that basal SEMA3G levels are unaffected by presence of tau, amyloid or alpha-synuclein pathology.

**Figure 6:**
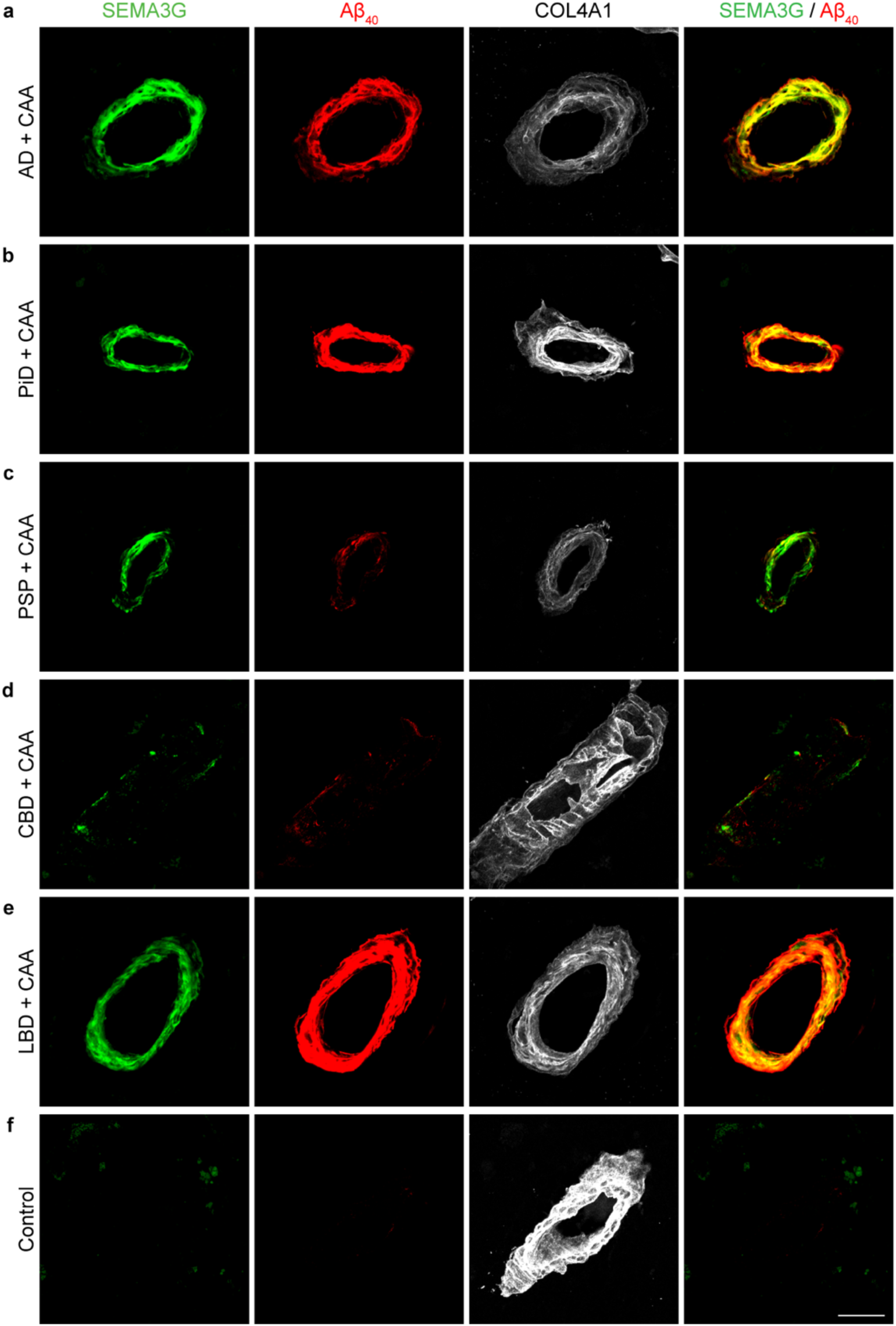
SEMA3G is a selective marker of CAA in a variety of neurodegenerative diseases. **a,** Representative image of SEMA3G accumulation in CAA^+^ vessels of AD + CAA cases with Aβ and tau pathology (*n* = 6). **b-d,** Representative images of SEMA3G accumulation in CAA^+^ vessels of primary tauopathy cases; PiD + CAA (*n* = 2) **(b)**, PSP + CAA (*n* = 4) **(c)**, CBD + CAA (*n* = 1) **(d)**. **e,** Representative image of SEMA3G accumulation in CAA^+^ vessels of LBD + CAA cases with alpha-synuclein pathology (*n* = 2). **f,** Representative image of control cases (*n* = 5) showing no SEMA3G or Aβ40 signal. Scale bar = 25µm.

### SEMA3G delays the formation of Aβ40 and Aβ42, and binds to fibrils in vitro

Having observed strong SEMA3G and Aβ_40_ immunofluorescence in CAA^+^ vessels, and the strong PLA observed in cases with high CAA load, we next investigated the ability of SEMA3G to influence Aβ fibril formation. Thioflavin-T (ThT) assays were performed incubating Aβ_40_ or Aβ_42_ with SEMA3G at a 1:0.5, 1:0.1 or 1:0.05 molar ratio (*n* = 3-6 per condition) or with the control protein hen egg white lysozyme (HEWL) at a 1:0.5 molar ratio (*n* = 2 Aβ_40_, *n* = 3 Aβ_42_). SEMA3G had a pronounced effect on t_1/2_ of both Aβ_40_ and Aβ_42_ fibril formation at all molar ratios (Fig. 7). Delayed Aβ_40_ fibril formation was consistent across all ratios (Fig. 7a-b), while the effect on Aβ_42_ appeared dose dependant, approaching saturation at a ratio of 1:0.1 (Fig. 7c-d).

**Figure 7:**
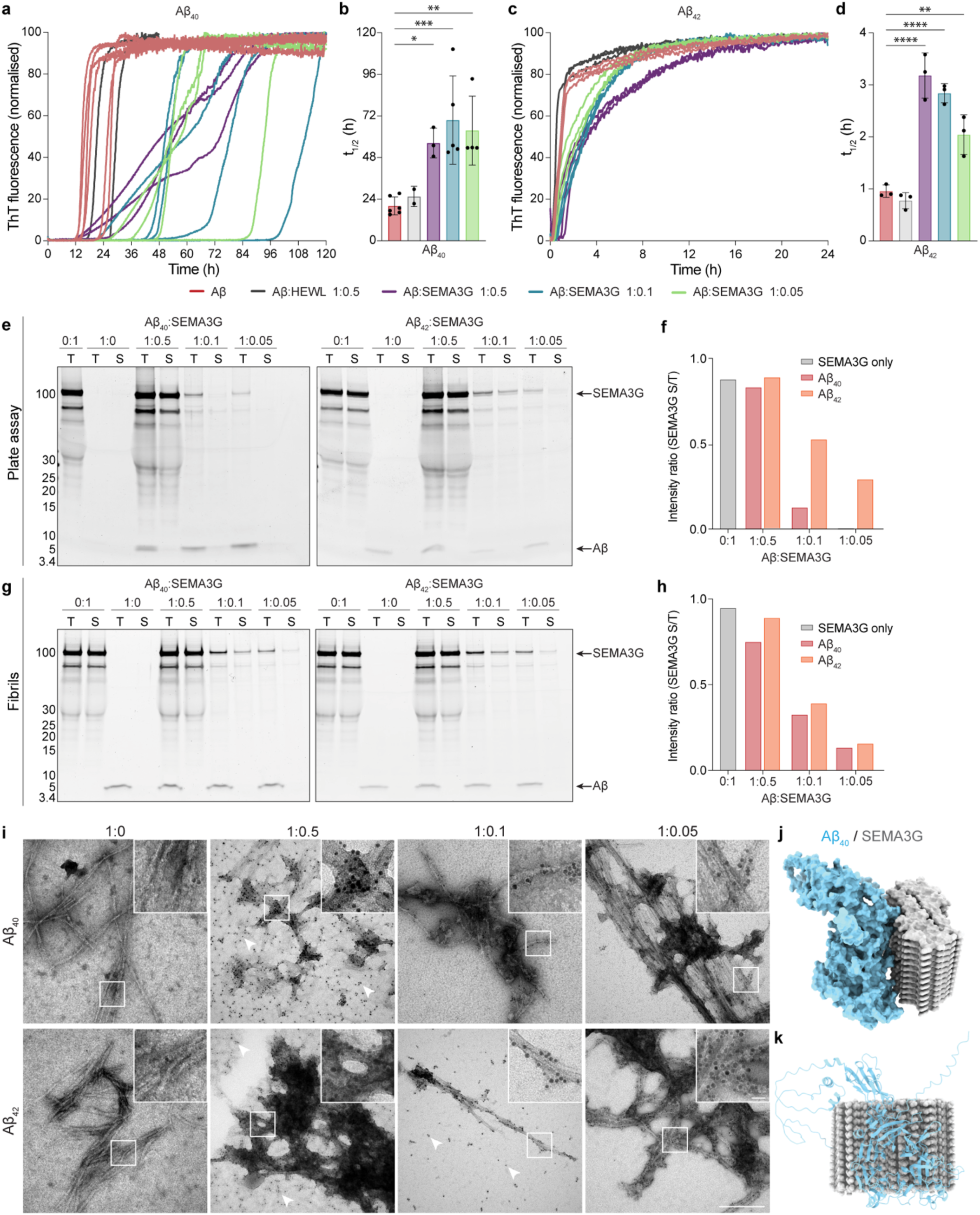
SEMA3G binds to Aβ40 and Aβ42 *in vitro.* **a-b,** Normalized thioflavin T (ThT) traces of 5µM Aβ40 assembled with a 1:0.5-0.05 ratio of SEMA3G or hen egg white lysozyme (HEWL; protein control) **(a),** showing increased t1/2 in the presence of SEMA3G **(b)**. **c-d,** Normalized ThT traces of 5µM Aβ42 assembled with a 1:0.5-0.05 ratio of SEMA3G or HEWL **(c)** showing increased t1/2 in the presence of SEMA3G **(d)**. Data show mean ± SD of *n* = 6 Aβ40, *n* = 2 Aβ40:HEWL 1:0.5, *n* = 3 Aβ40:SEMA3G 1:0.5, *n* = 5 Aβ40:SEMA3G 1:0.1, *n* = 4 Aβ40:SEMA3G 1:0.05, *n* = 3 all Aβ42 conditions; * *p* < 0.05, \*\**p* < 0.01, \*\*\**p* < 0.001, \*\*\*\**p* < 0.0001 determined by one-way ordinary ANOVA with Dunnett’s multiple comparisons test. **e,** SDS-PAGE gels showing total samples (T) and soluble fractions (S) following ThT assays. **f,** Intensity ratio of the SEMA3G band in soluble:total fractions. **g,** SDS-PAGE gels showing total samples (T) and soluble fractions (S) following incubation with preformed Aβ fibrils. **h,** Intensity ratio of the SEMA3G band in soluble:total fractions from preformed fibril incubation. **i,** Representative TEM images of samples from ThT assay plates shows SEMA3G coating fibrils of both Aβ40 and Aβ42 at all concentrations. Unbound SEMA3G was present in 1:0.5 Aβ40:SEMA3G, 1:0.5 Aβ42:SEMA3G and 0.1 Aβ42:SEMA3G conditions (arrows). Ni-NTA nanogold (black dots) indicates the presence of a single SEMA3G monomer. Scale bars = 200 nm, 20 nm (insets). **j,** Molecular docking illustrates relative size of SEMA3G molecule and Aβ40 fibril, represented as two 12-mer protofilaments. Docking predicts potential interaction between SEMA3G and the face of the Aβ40 fibrils. **k,** The face of the β-propeller domain of SEMA3G is oriented against the fibril surface.

To assess whether SEMA3G was bound to the resulting Aβ fibrils, ThT assay end-point samples were centrifuged to pellet Aβ fibrils and oligomers and analysed by SDS-PAGE. Additionally, to investigate whether SEMA3G would bind to preformed Aβ fibrils, equal concentrations of SEMA3G or HEWL control (*n* = 2 per condition) were incubated with preformed fibrils of either Aβ_40_ or Aβ_42_, centrifuged and the depletion of Aβ peptide and SEMA3G was monitored. Aβ fibrils were expected to pellet completely under these conditions, and SEMA3G was expected to remain in the soluble fraction if unbound.

SEMA3G was pulled down with both Aβ_40_ and Aβ_42_ in both conditions (Fig. 7e-h), while HEWL control remained in solution (Extended Data Fig. 7). SEMA3G was most effectively depleted from the soluble fraction in Aβ_40_ assay plate samples, where a complete shift from soluble to insoluble was observed at 1:0.05 ratio, and a near complete shift was observed at 1:0.1 (Fig. 7e-f). Depletion of SEMA3G from the soluble fraction was similarly observed in Aβ_42_ plate conditions, however this was to a lesser extent than Aβ_40_. Interestingly, soluble SEMA3G was consistently depleted when incubated with either Aβ_40_ or Aβ_42_ preformed fibrils (Fig. 7g-h).

We were next interested to determine if SEMA3G directly interacted with Aβ fibrils and if this interaction influenced fibril morphology. Transmission electron microscopy (TEM) with Ni-NTA nanogold staining was used to determine whether His-tagged SEMA3G was colocalized with the Aβ fibrils formed during the ThT assay. Large clusters of amorphous protein as well as smaller protein clusters, both positive for nanogold, were observed associated with Aβ_40_ and Aβ_42_ fibrils at all SEMA3G concentrations (Fig 7i). Background nanogold binding to free SEMA3G increased proportionally with SEMA3G concentration and was consistent with centrifugation assay data, with little unbound SEMA3G observed at 1:0.1 ratio with Aβ_40,_ and slightly more unbound SEMA3G observed at 1:0.1 with Aβ_42_ (Fig. 7i, arrows).

Having observed a preferential interaction of SEMA3G with Aβ_40_ *in vitro*, we next used *in silico* modelling to investigate potential interactions between SEMA3G and Aβ_40_ fibrils. The RosettaDock 5.0 protocol was utilized to dock the predicted three-dimensional structure of SEMA3G (AlphaFold2) onto the structure of amyloid filaments composed of Aβ_40_, as determined from the leptomeninges of individuals with AD and CAA ^27^. A model consisting of two 12-mer protofilaments was prepared. The docking revealed that if bound, one molecule of SEMA3G occupies a surface area of the amyloid fibril composed of ∼20 Aβ_40_ molecules (Fig. 7j). For each successful docking, multiple models were identified with CAPRI scores of 2 or 3 and low interface energies. The structure with the lowest interaction energy score predicted an interaction of SEMA3G with the face of the Aβ_40_ fibril and exhibited multiple sites of close contact (Fig. 7k). For both fibril face and end starting poses, only those with the SEMA3G β-propeller positioned so that loops containing residues 319, 448 and 554 were oriented towards the fibril surface produced valid predictions.

## Discussion

Our results show that SEMA3G is a highly specific marker of CAA in the brain. SEMA3G consistently accumulated in CAA^+^ vessels in both the leptomeninges and parenchyma in multiple brain regions, in a variety of neurodegenerative diseases and across the spectrum of CAA severity. The selective colocalization of SEMA3G with CAA and not amyloid plaques was consistently observed in all cases examined. We showed that SEMA3G was found in closer proximity to Aβ_40_, pyroglutamated Aβ and phosphorylated Aβ than with Aβ_42_ and that SEMA3G significantly slowed the aggregation of both Aβ_40_ and Aβ_42_ *in vitro*, with this effect being more pronounced for Aβ_40_.

The selective association of SEMA3G with CAA and not amyloid plaques is significant given that the vast majority of Aβ-interacting proteins accumulate in both CAA and amyloid plaques^15,19^. The SEMA3G selectivity for CAA was most clearly demonstrated in dyshoric plaques, where SEMA3G was abundant only in the vascular component and not in the neighbouring halo of parenchymal Aβ. This suggests that these two pools of Aβ are distinct and bind to different proteins, rather than reflecting vascular wall destruction resulting in Aβ leakage into the parenchyma, as has been previously hypothesized^2^. The selectivity of SEMA3G for CAA could be a consequence of the unique production of SEMA3G by arterial endothelial cells^16^, which limits the opportunity for SEMA3G to interact with Aβ outside of blood vessels. However, this simple explanation does not account for the fact that SEMA3G is a secreted protein demonstrated to exert paracrine effects on multiple cell types^20,21,28^, including neurons in AD affected brain regions^22^. This suggests that SEMA3G has the opportunity to interact with Aβ in amyloid plaques and selectively does not. A second possible explanation is that SEMA3G selectively interacts only with specific types or conformations of Aβ that are present in CAA and not amyloid plaques. Previous studies showing that vascular and parenchymal Aβ adopt different fibril conformations^27,29,30^ and are enriched in different Aβ proteoforms^31^ support this hypothesis. Our *in vitro* results showing a stronger interaction of SEMA3G with Aβ_40_ than Aβ_42_ also supports this hypothesis given the preferential deposition of Aβ_40_ in large CAA^+^ vessels in comparison to Aβ_42_ in amyloid plaques^32–36^. Our observation of clusters of amyloid plaques highly enriched in Aβ_40_ but negative for SEMA3G indicate that factors beyond Aβ_40_ deposition alone must underlie the selective accumulation of SEMA3G in CAA. Additionally, our PLA results show that SEMA3G had a close proximity to both pyroglutamated Aβ and phosphorylated Aβ in CAA, both of which are also abundant in amyloid plaques in addition to CAA^31,37–41^. Therefore, while the strong interaction between SEMA3G with Aβ_40_ could partially account for preferential accumulation of SEMA3G in CAA, more research is needed to determine why SEMA3G only accumulates in CAA and not amyloid plaques.

Another notable observation was the widespread accumulation of SEMA3G throughout CAA^+^ blood vessels, extending beyond regions of Aβ deposition. This was particularly evident in vessels with mild CAA, where small Aβ puncta were associated with widespread SEMA3G accumulation. This suggests that early SEMA3G accumulation in CAA^+^ vessels is driven by factors beyond direct interaction with Aβ. In other contexts, SEMA3G appears to play a protective role in response to vascular injury. Hypoxic or ischemic stress increases SEMA3G expression, where it promotes endothelial junction stability and limits excessive angiogenesis during vascular remodelling^23,24^. SEMA3G is also upregulated after arterial injury, where it promotes smooth muscle cell proliferation and migration to support vessel wall repair^21^. Given these established protective roles, we hypothesize that SEMA3G is produced by arterial endothelial cells as an early protective response to vascular injury during CAA development. Consistent with these previous studies, increased SEMA3G expression may reflect an attempt to promote the recruitment or proliferation of vascular smooth muscle cells to compensate for those damaged by Aβ deposition. Alternatively, it may reflect a response to local hypoxic or ischemic stress resulting from CAA-associated vascular dysfunction. As CAA progresses, our results suggest that SEMA3G increasingly interacts with Aβ in the vessel walls. While this this sequestration of SEMA3G may compromise its protective actions for vessel wall repair, it also drives a secondary protective mechanism of slowing Aβ_40_ aggregation, as evidenced by our *in vitro* studies. One downside of the SEMA3G sequestration in CAA is that it could potentially contribute to neuronal dysfunction, given that SEMA3G also directly regulates synaptic plasticity and hippocampal-dependent memory^22^. Together, this suggests that SEMA3G likely functions as a protective protein in CAA^+^ vessels but could inadvertently contribute to neuronal dysfunction as a consequence of sequestration into CAA.

While only a small number of cases in our cohort had capillary CAA, it was clear from these cases that SEMA3G did not colocalize with capillary CAA, suggesting that SEMA3G is specific to arterial and arteriole CAA. The lack of SEMA3G accumulation in capillary CAA could be due to the increased deposition of Aβ_42_ rather than Aβ_40_ in capillary CAA^42,43^. Alternatively, this could be due to the low expression of SEMA3G by capillary endothelial cells ^16,17^ or the lack of smooth muscle cell layer in capillaries, which is the primary site of SEMA3G expression in arteries and arterioles.

The high specificity of SEMA3G for CAA^+^ vessels in both the parenchyma and leptomeninges across all diseases and brain regions examined suggests that SEMA3G may be a promising biomarker for CAA, either as a fluid biomarker or PET tracer. While previous proteomic studies of CSF or plasma have not yet identified SEMA3G as a significant biomarker of CAA, this is potentially because most prior studies have not examined SEMA3G biofluid changes in individuals with neuropathologically confirmed CAA^44,45^. While one previous study did assess proteomic differences in the CSF of neuropathologically confirmed sporadic CAA cases, SEMA3G levels were too low abundance (detected in less than 15% of samples) to draw definitive conclusions about its biomarker potential at this stage^46^. Therefore, future studies using alternative approaches to assess the potential of SEMA3G as a biofluid biomarker or PET tracer of CAA are warranted.

In conclusion SEMA3G is a novel, highly specific marker of CAA in the brain. The early, selective accumulation of SEMA3G in CAA^+^ vessels suggests that it may have an important mechanistic role in CAA development. Future studies examining the functional role of SEMA3G in CAA development and its potential as a fluid biomarker or PET tracer for CAA are recommended.

## Methods

### Human Tissue Samples

Human brain tissue was acquired from Rush University (USA), the New York University Alzheimer’s Disease Center (USA), and the Sydney Brain Bank (Australia), which all provide human brain tissue from ethically approved longitudinally assessed regional brain donor programs on neurodegenerative diseases. Samples were obtained from multiple brain banks based on convenience and availability. Brain tissue was acquired under protocols with Institutional Review Board (IRB) approval at NYU Grossman School of Medicine, Rush University and the Southeastern Sydney and Illawarra Local Health District and the Universities of New South Wales and Sydney, Australia. In all cases, written informed consent for research was obtained from the patient or legal guardian, and the material used had appropriate ethical approval for use in this project. All patients’ data and samples were coded and handled according to NIH and NHMRC guidelines to protect patients’ identities.

Inferior temporal cortex and primary visual cortex tissue was obtained from Rush University, which was from cases part of the Religious Orders Study (ROS) and Memory and Aging Project (MAP) cohorts^47^. Clinical assessments and neuropathology were performed at Rush University^4,48,49^. Cases were initially stratified by neuropathological ABC and CAA scores. Cases were then screened using total Aβ immunohistochemistry and Aβ plaque and CAA load quantified. The following inclusion criteria was then used to stratify cases for SEMA3G investigation: CAA and plaque load < 0.5% for control; CAA load < 0.5, plaque load > 0.5% for CAA^−^ P^+^; CAA load 0.5-1.4%, plaque load > 0.5% for mild CAA; CAA load > 1.4%, plaque load > 0.5% for moderate CAA; CAA load > 0.5%, plaque load < 0.5% for CAA^+^ P^−^. Cases were prioritized to exclude those with high TDP-43 and Lewy body pathology. Tissue included in this study from this cohort included 8 mm formalin-fixed paraffin-embedded (FFPE) sections from the inferior temporal cortex of control (*n* = 20), mild CAA (*n* =14), moderate CAA (*n* = 16), CAA^+^ P^−^ (*n* = 5), and CAA^−^ P^+^ cases (*n* = 20). A smaller primary visual cortex cohort was selected comprising control (*n* = 5), mild CAA (*n* = 6), moderate CAA (*n* = 4), CAA^+^ P^−^ (*n* = 4), and CAA^−^ P^+^ cases (*n* = 4). All groups were balanced for sex and postmortem interval where possible. Superior frontal cortex tissue was obtained from the Sydney Brain Bank (Australia). Cases were screened for the presence of Aβ plaques or CAA using pan-Aβ immunohistochemistry. Any cases exhibiting CAA were deemed CAA^+^, regardless of severity. This included 8 mm FFPE sections from *n* = 5 control, *n* = 6 AD CAA^+^, *n* = 4 AD CAA^−^, *n* = 1 CBD CAA^+^, *n* = 5 CBD CAA^−^, *n* = 2 PiD CAA^+^, *n* = 5 PiD CAA^−^, *n* = 4 PSP CAA^+^, *n* = 5 PSP CAA^−^, *n =* 2 LBD CAA^+^, and *n* = 5 LBD CAA^−^ cases. Additional primary visual cortex tissue was obtained from New York University Alzheimer’s Disease Center (USA). This tissue included 8 μm FFPE sections from *n* = 3 AD CAA^+^ cases. Case-specific details for all cases are summarized in Supplementary Table 1.

### Immunohistochemistry

FFPE tissue sections underwent fluorescent immunohistochemistry using the method described in^50^ with some modifications. Briefly, sections were deparaffinized and rehydrated through a series of xylene and ethanol washes. Antigen retrieval was achieved using 99% formic acid for 7 min followed by boiling in citrate buffer for 21 min (0.05 mM sodium citrate, 0.05% Tween-20, pH 6) and incubation in proteinase K for 5 min at RT (Sigma, P5568, 0.125mg/mL in PBS). Sections were blocked in 10% normal horse serum and incubated with primary antibodies in 4% normal horse serum overnight at 4°C (Supplementary Table 2). The specificity of the anti-SEMA3G antibody was confirmed by preabsorption with SEMA3G recombinant protein (Abclonal RP01400) prior to immunostaining (Supplementary Figure 1). Fluorophore conjugated secondary antibodies and Hoechst 33342 (Sigma, B2261, 1:2000) were applied for 2 h at room temperature. For Aβ proteoform and neurodegenerative disease cohort stains, sections were incubated with TrueBlack Autofluorescence Quencher (Cell Signaling Technology, 92401, 1:20 in 70% ethanol) prior to coverslipping with Antifade ProLong Glass (Invitrogen, P36984). Whole slide images were acquired using an Olympus VS200 Slide Scanner at 10x (NA 0.4) magnification. Representative 63x images (NA 1.4) were captured on a Leica Stellaris 8 Confocal microscope. For Figures 4 and 5, representative images were taken from the same vessel on serial sections.

### Immunohistochemistry Analysis

SEMA3G load in blood vessels, CAA, Aβ plaques and grey matter was assessed in inferior temporal and primary visual cortex sections. Grey matter and meningeal regions were manually annotated in QuPath (v0.5.0). Pixel classifiers were trained in QuPath to recognize blood vessels and amyloid plaques, based on COL4A1 and Aβ channels respectively. Classifiers were used to generate masks of blood vessels and Aβ in each section. To minimize autofluorescence detection, positive signal was restricted to >40 µm^2^ and >200 µm^2^ respectively. Images with corresponding masks were then exported as .tif images to ImageJ2 (1.54p). Areas of CAA in the grey matter were identified by finding overlapping positive areas in blood vessel and Aβ masks. CAA detections were restricted to >40 µm^2^ to minimize inclusion of areas in which amyloid plaques encroached on blood vessels. Blood vessels containing CAA were identified and separated out as CAA^+^ vessels (Extended Data Fig. 1). Meningeal CAA was identified by subtracting empty channel 532 from Aβ, thresholding and masking positive signal. Empty channel 532 was then subtracted from the SEMA3G channel to minimize autofluorescent signal, a threshold determined based on the negative control, and signal measured within each region of interest.

### Statistical Analysis

Analysis was performed in GraphPad Prism (v10.0.3). Each dataset was assessed for Gaussian distribution using a Shapiro-Wilk normality test. Normally distributed datasets were analyzed using ordinary two-way ANOVA with Tukey’s multiple comparisons test, Pearson correlation or Welch’s t-test. Non-normal datasets were analyzed using Kruskal-Wallis test with Dunn’s post-hoc, nonparametric Spearman correlation or Mann-Whitney two-tailed U test where applicable.

### Proximity Ligation Assay (PLA)

Serial sections from *n* = 3 primary visual cortex FFPE tissue sections underwent deparaffinization, rehydration, antigen retrieval and primary antibody steps as above. The secondary antibody for COL4A1 and Hoescht 33342 (Sigma, B2261, 1:2000) were applied in solution with DuoLink PLUS (Sigma, DUO92002, 1:5) and MINUS (Sigma, DUO92004, 1:5) probes for 1 hr at 37°C. Ligation and amplification (Sigma, DUO92013) were performed as per the manufacturer protocol prior to coverslipping with Antifade ProLong Glass (Invitrogen, P36984). Whole slide images were acquired using an Olympus VS200 Slide Scanner at 20x (NA 0.8) magnification. Representative 63x images (NA 1.4) were captured on a Leica Stellaris 8 Confocal microscope.

### Protein stock preparation for in vitro characterisation

Aβ_40_ (rPeptide, A11532) and Aβ_42_ (rPeptide, A11632) lyophilised powder was dissolved in ice cold 50 mM NaOH to 1 mg/mL concentration. Stocks were bath sonicated on ice for 10 min then centrifuged through Ultrafree® 0.22 µm PVDF filter (Merck), snap frozen and stored at −80°C until use. SEMA3G (Abclonal, RP01400) was dissolved in ddH_2_O to a concentration of 0.5 mg/mL, then centrifuged at 16 000 × *g* for 10 minutes at 4°C to remove large aggregates. SEMA3G concentration was confirmed by 280 nm absorbance using Beer-Lambert’s law with ε = 106 685cm^−1^ M^−1^.

### Thioflavin T assays

5 µM Aβ and 2.5–0.25 µM SEMA3G or 2.5 µM lysozyme were diluted into half-area, non-binding, 96-well black clear-bottom assay plates (Corning, 3881) containing 50% 100 mM phosphate pH 6.5, 50% 25 mM NaH_2_PO_4_ pH 7.4, 150 mM NaCl and 40 µM thioflavin T (ThT). Aβ aliquots were thawed then bath sonicated on ice for 10 minutes just prior to addition to microplates. Fluorescence was measured using a BMG PolarStar microplate reader. Plates were shaken initially at 500 rpm for 30 seconds (double orbital) then left quiescent for 5 days at 37°C. ThT fluorescence emission at 480 nm was read every 5 minutes with excitation at 440 nm.

### Statistical Analysis

ThT assay analyses were conducted using GraphPad Prism (11.0.0). ThT data was collated and normalised by the maximum fluorescence value from each triplicate at 24 h (Aβ_42_) or 120 h (Aβ_40_). Datasets that showed no increase in fluorescence were excluded. Normalised data was fitted to a sigmoidal dose-response curve (Prism analysis for nonlinear regression (curve fit): [Agonist] vs. normalised response – variable slope) for estimation of t_1/2_ values for each dataset. Statistical analysis was performed using an ordinary one-way ANOVA with Dunnett’s multiple comparisons test.

### Aβ fibril formation

Aβ_40_ and Aβ_42_ aliquots were assembled at 20 μM in triplicate in a plate reader as described for ThT assay methods. Fibrils were recovered from wells, then wells were washed once with an equal volume of PBS for a final concentration of 10 μM. Fibrils were bath sonicated on ice for 5 min to disperse fibril clusters, then mixed 1:1 with PBS to a final concentration of 5 µM, or with SEMA3G to final molar ratios of 1:0.5, 1:0.1 or 1:0.05.

### Aβ fibril centrifugation

Samples from ThT assays or from fibril incubations were reserved as total samples. Aliquots from the same well were centrifuged at 100 000 × *g* for 30 min at 4 °C using an Optima^TM^ MAX-XP ultracentrifuge (Beckman Coulter), then an equal volume of supernatant was taken for SDS-PAGE analysis. Samples were prepared with 1X NuPAGE sample reducing agent (Invitrogen) and 1X Tricine SDS loading dye (Invitrogen) and loaded onto 10–20% tricine gels (Invitrogen) for electrophoresis in Tricine SDS running buffer (Invitrogen) at 125 V for 80 min. Gels were stained with Sypro^TM^ Ruby protein gel stain (Invitrogen) according to manufacturer’s protocol and imaged on a ChemiDoc MP Imaging System (BioRad).

### Gel intensity analysis

Image analysis was conducted using ImageJ (1.54p). Unsaturated images were utilised for analysis. ImageJ built-in tools for gel analysis were used to select SEMA3G bands across each gel and plot lane profiles. Baselines were set to remove background intensity, and the area under the resulting curve was measured. Area of SEMA3G bands from the soluble fraction was divided by area of SEMA3G bands from the total fraction, and ratios were plotted using GraphPad Prism (11.0.0).

### Transmission Electron Microscopy

Transmission electron microscopy was performed at the Sydney Microscopy and Microanalysis Core Research Facility. Carbon-coated copper 200 mesh grids coated with formvar support film (ProSciTech Pty Ltd.) were glow discharged at 20 mA for 30 s, then 5 μL of sample was dropped onto grid and incubated for 5 min at room temperature. Grids were washed three times with ddH_2_O then floated on a droplet of 30X diluted 10 nm Ni-NTA nanogold (Nanoprobes, 2084) in PB/PBS containing 1 μM BSA for 30 min at room temperature. Grids were washed once with ddH_2_O then stained with 2% uranyl acetate for 2 minutes and left to dry. Images were captured on a Tecnai T12 microscope at 120 kV and processed using DigitalMicrograph software (GATAN).

### Docking SEMA3G onto Aβ40 amyloid fibril structure

A 20-monomer protofilament was reconstructed from the model of Aβ_40_ (PDB8QNZ and EMDB18509) described in^27^. PyMol was used to position the AlphaFold model of SEMA3G (AF-Q9NS98-F1) in four different starting poses for docking onto the Aβ40 filaments: SEMA3G positioned at the short edge of the fibril, or against the fibril face, with either face of the seven-bladed β-propeller of the Sema domain positioned towards the fibril. Docking was performed using RosettaDock 5.0^51,52^, without ensemble docking mode. Docking of a single, fixed-backbone input structure for the fibril and SEMA3G was predicted by performing a Monte Carlo coarse-grain stage search, followed by an all-atom refinement stage to optimise rigid-body orientation and side-chain conformation. All decoy structures with CAPRI scores of >2 were assessed for area of interface. Chimera X^53^ was used for preparation of Fig. 7j-k.

## Data Availability

All data supporting the findings of this study are available from the corresponding author upon reasonable request.

## Acknowledgements

The authors acknowledge the technical and scientific assistance of Sydney Microscopy & Microanalysis, the University of Sydney node of Microscopy Australia. The authors also acknowledge and the patients and families for their generous brain donations to the Sydney Brain Bank (which is supported by Neuroscience Research Australia), the New York University Alzheimer’s Disease Centre, and Rush University Medical Centre.

## Author Contributions

E.D. conceived and designed this study. K.B. and A.S were responsible for immunohistochemistry. K.B was responsible for microscopy, analysis and figure generation. D.C. assisted with microscopy analysis. J.B. was responsible for Thioflavin-T assays and electron microscopy. E.D. and M.S. provided expert advice on the interpretation of data. T.W, J.S. and A.F. performed neuropathological characterization and provided human tissue. E.D. and K.B. wrote the manuscript with input from co-authors. All authors read and approved the final manuscript.

## Extended Data Figures

**Extended Data Figure 1:**
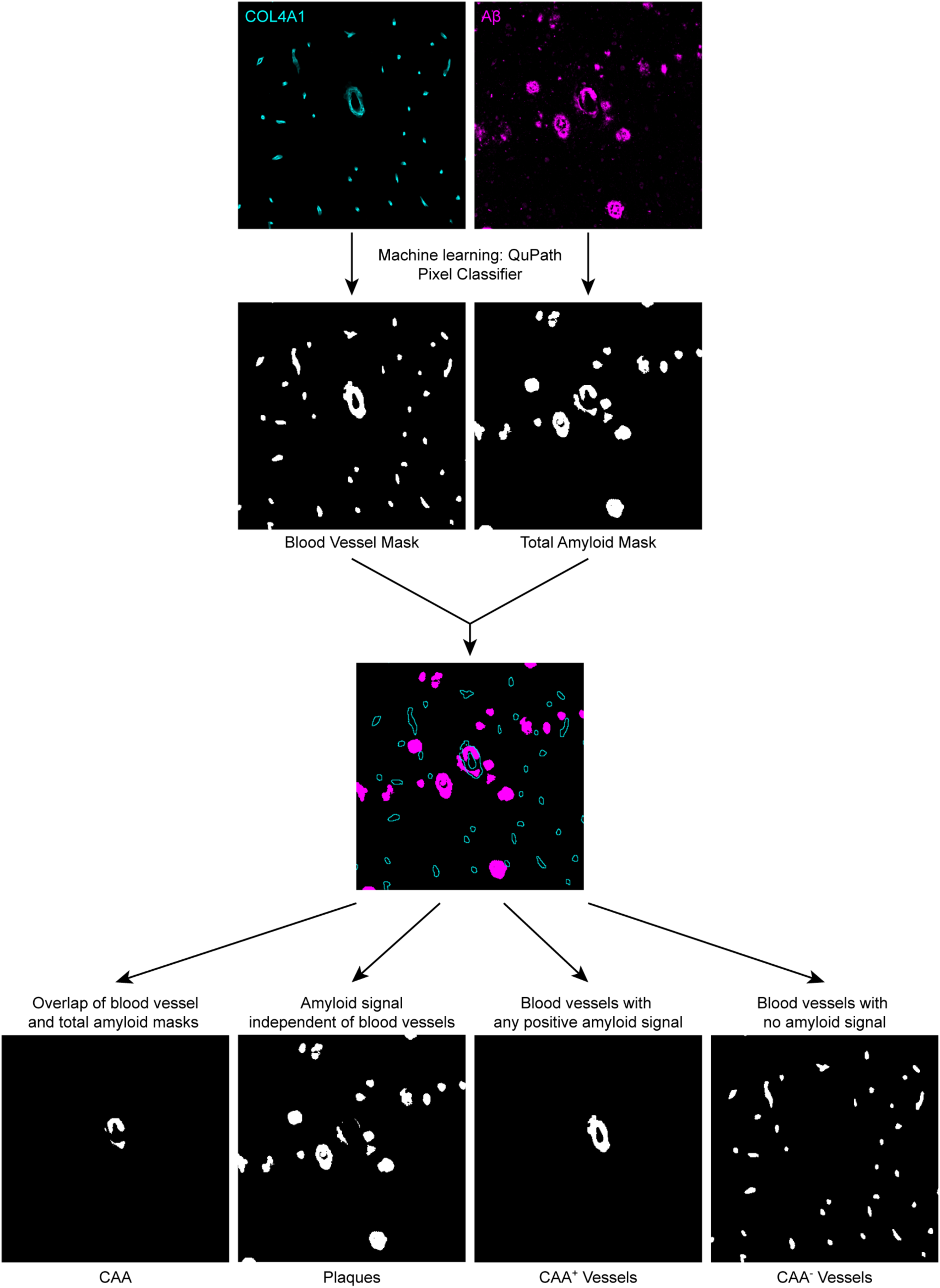
Classification of CAA, amyloid plaques, CAA^+^ and CAA^−^ blood vessels. Immunofluorescent signal from COL4A1 and pan Aβ were used to train pixel classifiers for blood vessels and total amyloid respectively. Classifier masks were then used to identify regions of CAA, amyloid plaques, CAA^+^ and CAA^−^ blood vessels.

**Extended Data Figure 2:**
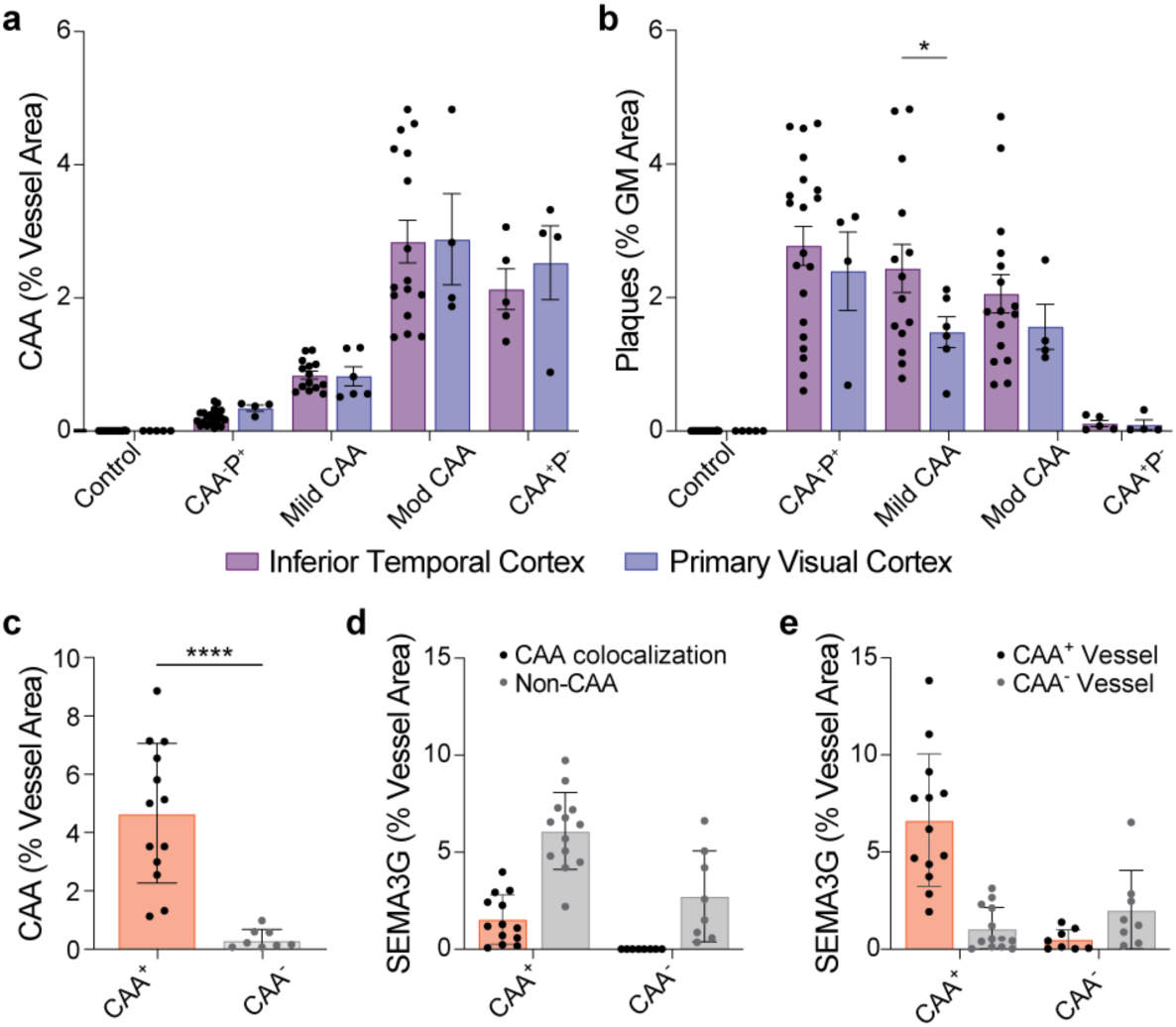
Characterization of inferior temporal and primary visual cortices. **a-b,** Quantification of CAA load **(a)** and plaque load **(b)** in the inferior temporal and primary visual cortices. Data show mean ± SD of *n* = 20 control, CAA^−^ P^+^, *n* = 14 Mild CAA, *n* = 16 Moderate CAA, *n* = 5 CAA^+^ P^−^ inferior temporal cortex cases; *n* = 5 control, *n* = 6 Mild CAA, *n* = 4 CAA^−^ P^+^, Moderate CAA, CAA^+^ P^−^ primary visual cortex cases. \**p*<0.05, determined by a two-way ANOVA test with Tukey’s post-hoc analysis. **c,** Classification of primary visual cortex cases into CAA^+^ or CAA^−^ based on leptomeningeal CAA load. \*\*\*\**p*<0.0001, determined by Welch’s t-test. **d,** Quantification of SEMA3G colocalization with Aβ in leptomeningeal vessels. **e,** Quantification of SEMA3G localization to CAA^+^ and CAA^−^ vessels in leptomeningeal vessels. Data in (c-e) show mean ± SD of *n* = 13 CAA^+^, *n* = 8 CAA^−^.

**Extended Data Figure 3:**
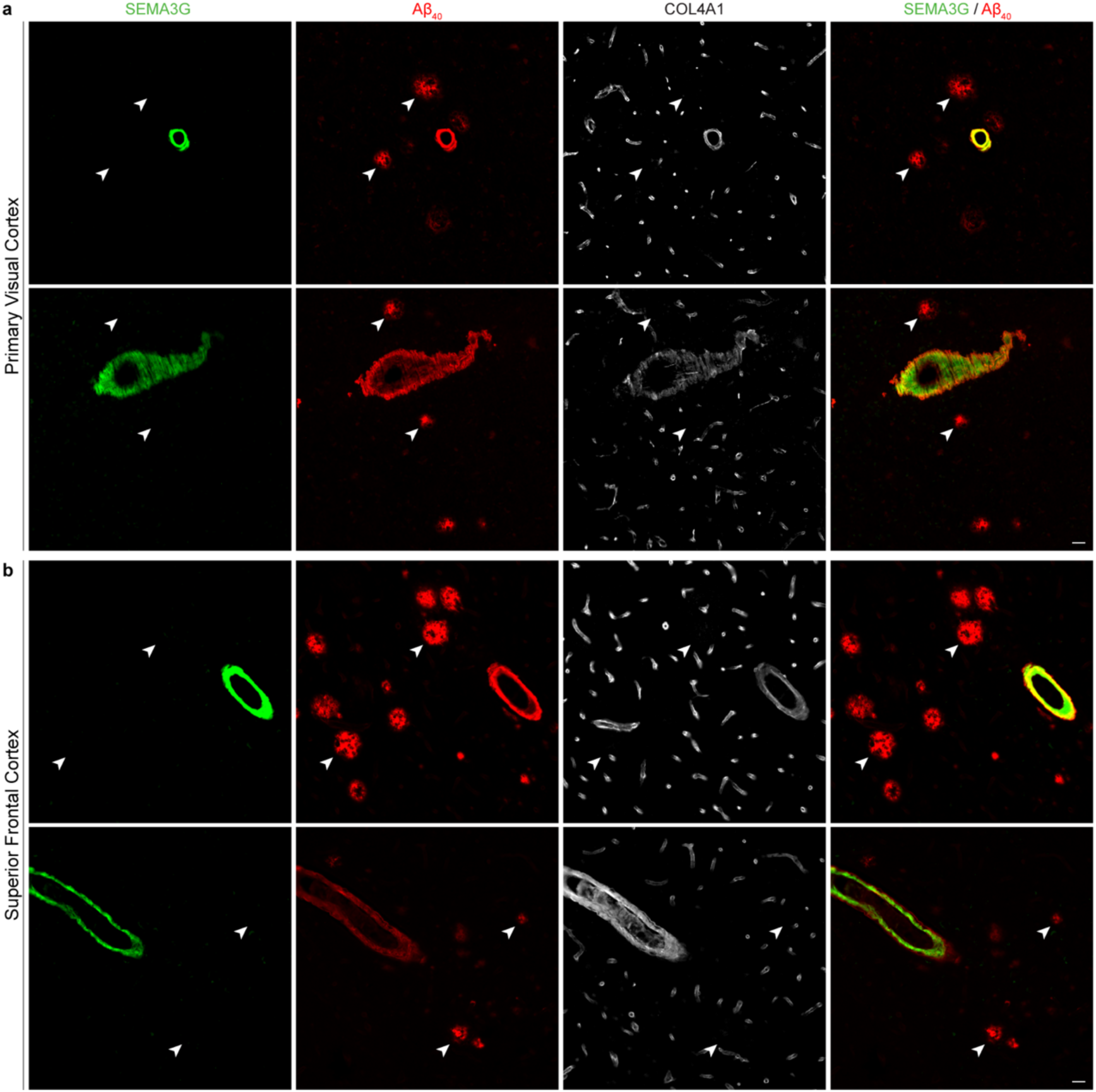
SEMA3G is not observed in Aβ40-positive amyloid plaques. **a-b,** Aβ40-positive amyloid plaques were negative for SEMA3G (arrows) in both the primary visual **(a)** and superior frontal **(b)** cortices, despite strong SEMA3G signal evident in Aβ40-positive CAA in nearby vessels. Scale bars = 25 µm.

**Extended Data Figure 4:**
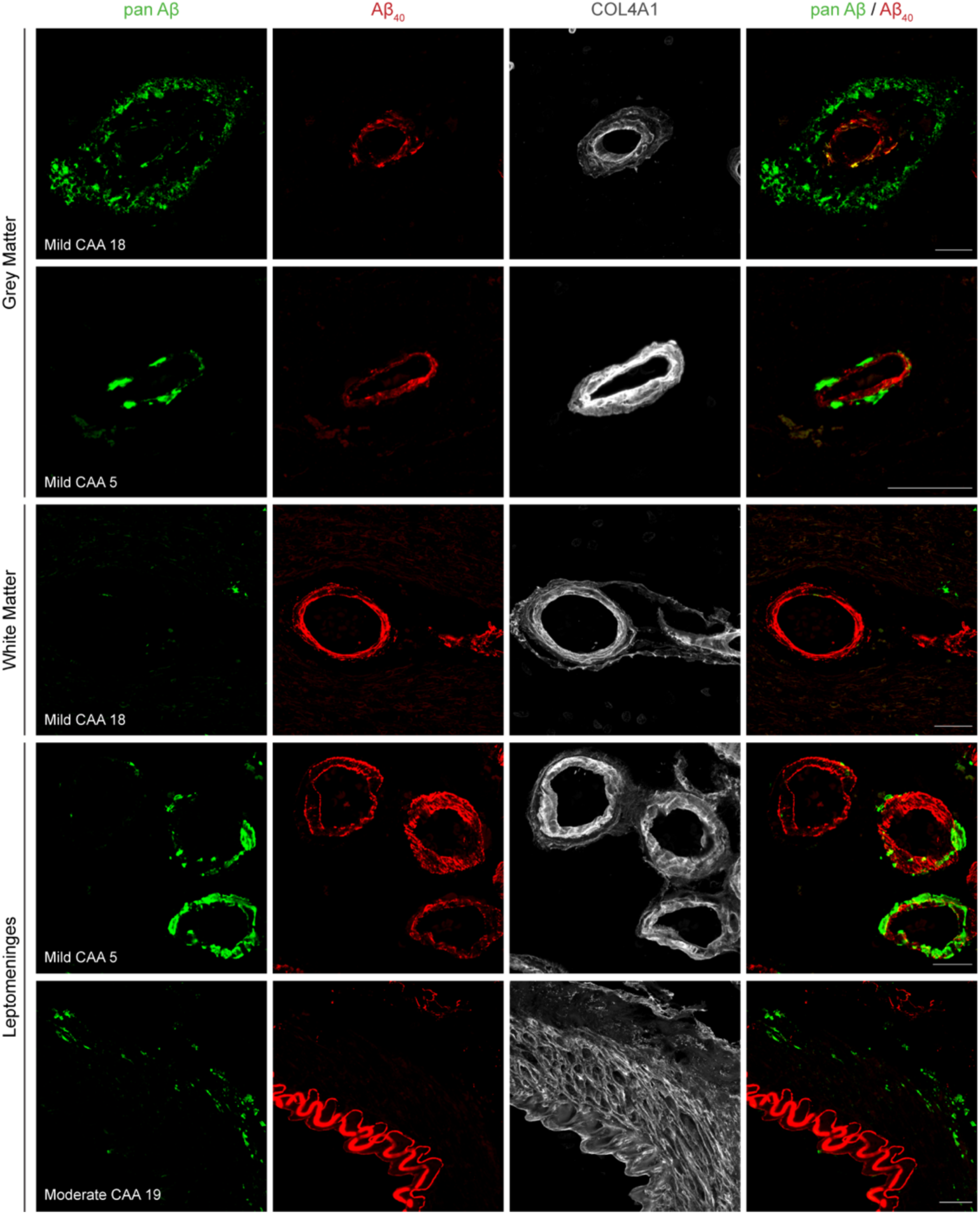
Immunostaining for Aβ40 detects CAA more extensively than a pan-Aβ antibody. Representative images showing different staining patterns of CAA using Aβ40 and pan-Aβ antibodies. Aβ40 immunostaining consistently revealed more extensive CAA than the pan-Aβ antibody across a range of CAA severities in the grey matter, white matter and leptomeninges. Scale bars = 25µm

**Extended Data Figure 5:**
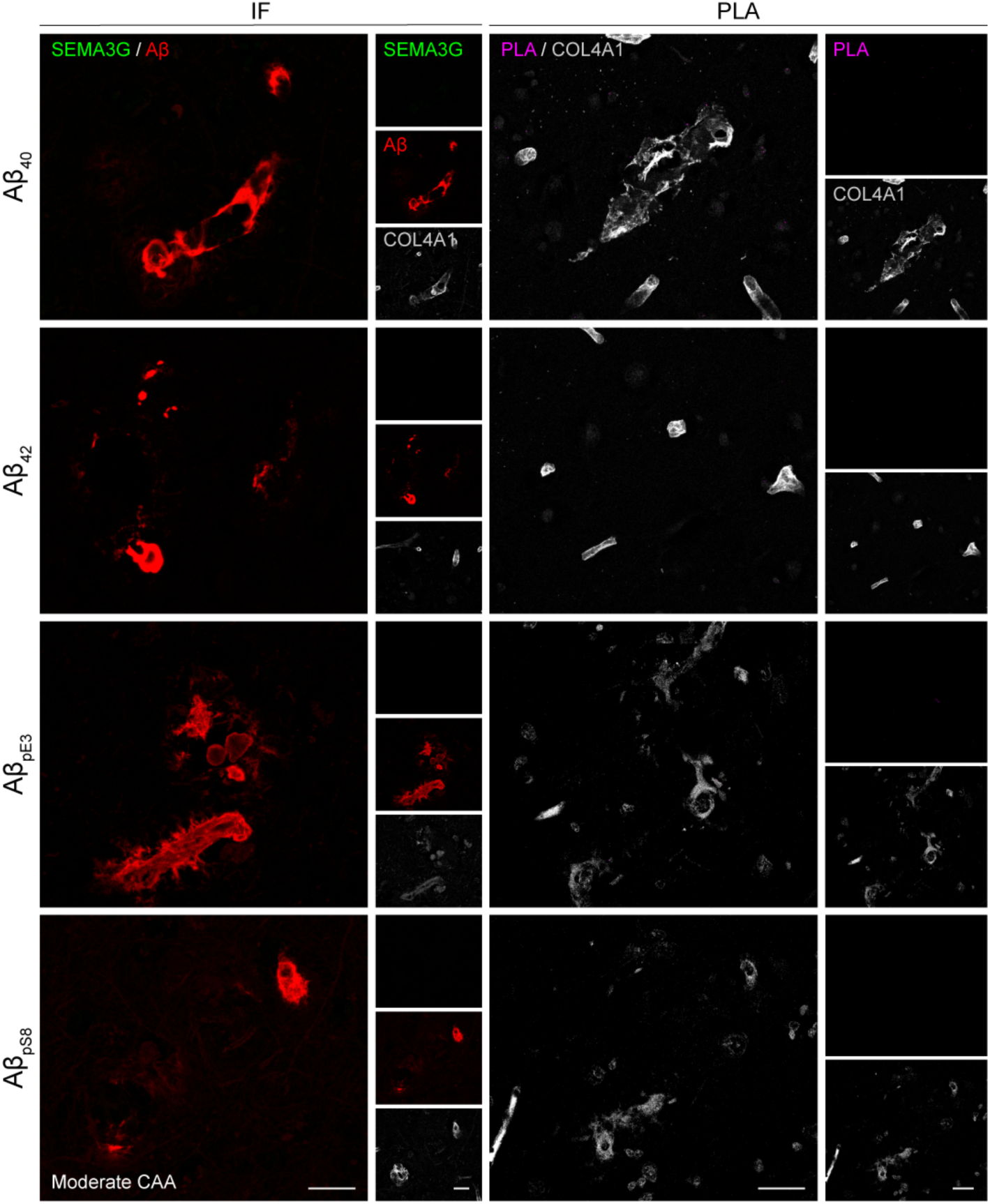
SEMA3G is not detected in capillary CAA. Capillary CAA positive for Aβ40, Aβ42, AβpE3 and AβpS8 was negative for SEMA3G immunofluorescence. PLA also did not reveal close proximity of SEMA3G with any Aβ proteoform in capillaries with CAA. Scale bars = 25 µm.

**Extended Data Figure 6:**
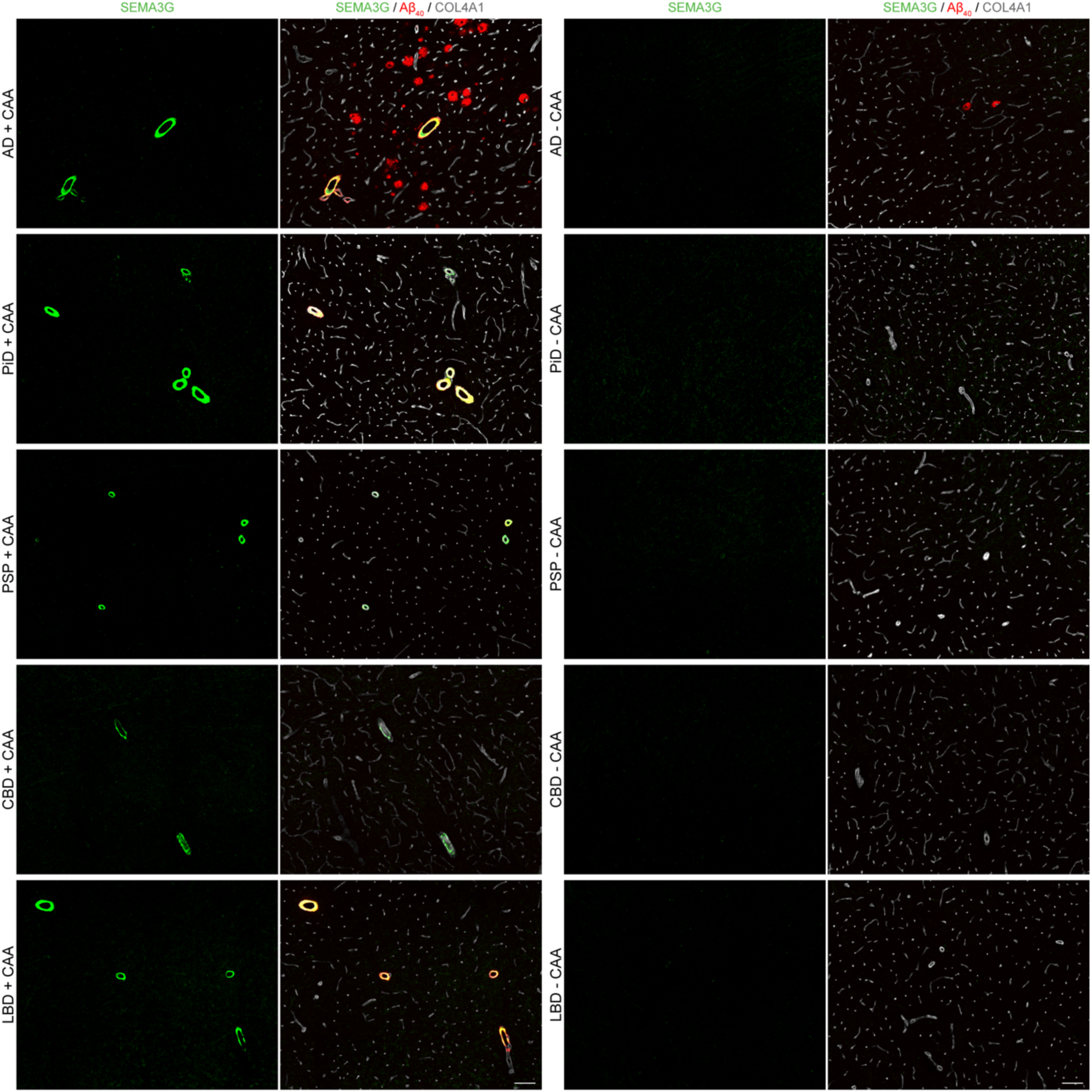
SEMA3G is highly specific to CAA^+^ vessels in a range of neurodegenerative diseases. Representative images showing SEMA3G immunofluorescence only in CAA^+^ vessels of neurodegenerative disease cases. SEMA3G was absent in amyloid plaques of AD + CAA cases. Minimal SEMA3G signal was detected in CAA-negative cases, regardless of neurodegenerative pathology. Images are representative of *n* = 6 AD + CAA, *n* = 2 PiD + CAA, *n* = 4 PSP + CAA, *n* = 1 CBD + CAA, *n* = 2 LBD + CAA, *n* = 4 AD - CAA, *n* = 5 PiD - CAA, PSP - CAA, CBD - CAA, LBD - CAA. Scale bars = 100 µm.

**Extended Data Figure 7:**
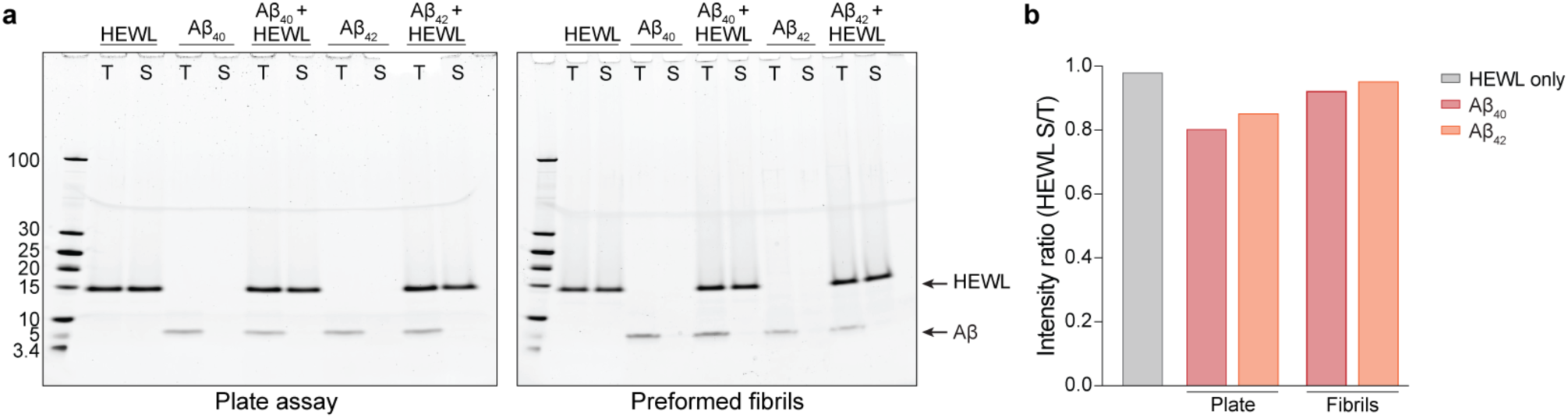
Hen egg white lysozyme (HEWL) control for Thioflavin T assays. **a,** SDS-PAGE gels showing total samples (T) and soluble fractions (S) following ThT assays or preformed fibril incubation. **b,** Intensity ratio of the HEWL band in soluble:total fractions, showing no depletion in any condition.

