## Supplementary Figure 1 for "Semaphorin 3G (SEMA3G) is a highly selective marker of cerebral amyloid angiopathy"

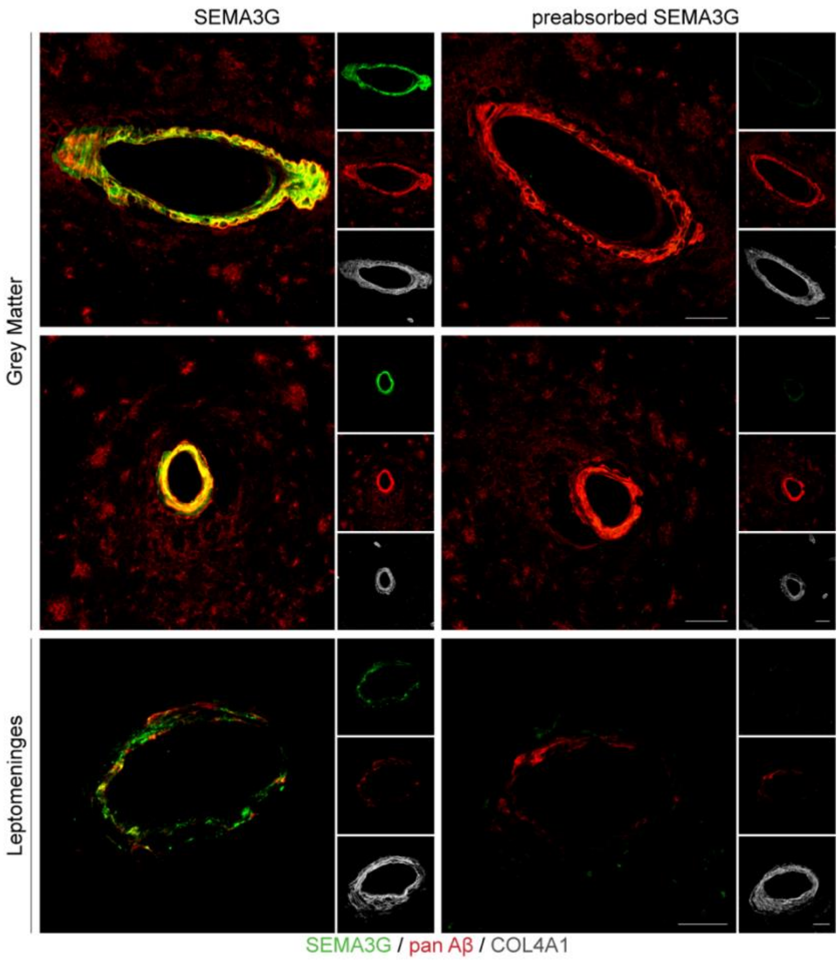


Supplementary Data 3: Validation of SEMA3G antibody HPA001761
SEMA3G antibody was preabsorbed by adding 5x excess SEMA3G protein (Abclonal, RP01400) by weight and incubating for 1 hr at RT with gentle shaking. The preabsorbed antibody was then used in place of a regular primary antibody for immunohistochemistry alongside a positive control. Absence of immunofluorescence with the use of preabsorbed antibody confirms specificity of the antibody for SEMA3G. Scale bars = 25 μm.
